# Text-related functionality and dynamics of visual human pre-frontal activations revealed through neural network convergence

**DOI:** 10.1101/2024.04.02.587774

**Authors:** Adva Shoham, Rotem Broday-Dvir, Itay Yaron, Galit Yovel, Rafael Malach

## Abstract

The functional role of visual activations of human pre-frontal cortex remains a deeply debated question. Its significance extends to fundamental issues of functional localization and global theories of consciousness. Here we addressed this question by comparing, dynamically, the potential parallels between the relational structure of prefrontal visual activations and visual and textual-trained deep neural networks (DNNs). The frontal visual relational structures were revealed in intra-cranial recordings of human patients, conducted for clinical purposes, while the patients viewed familiar images of faces and places. Our results reveal that visual relational structures in frontal cortex were, surprisingly, predicted by text and not visual DNNs. Importantly, the temporal dynamics of these correlations showed striking differences, with a rapid decline over time for the visual component, but persistent dynamics including a significant image offset response for the text component. The results point to a dynamic text-related function of visual prefrontal responses in the human brain.

## Introduction

While a large body of human brain research has centered on characterizing the functional organization of the visual system located at the posterior part of the brain^1^, a consistent finding in visual research highlights visual pre-frontal activations as well. These activations, although showing substantial report-related modulations (e.g.,^2^), have been demonstrated also in relatively passive report-free situations such as movie watching ^3–7^ or viewing static stimuli^8–11^. Furthermore, there is an ongoing debate concerning the functional role of these pre-frontal visual activations in human mental processes and behavior. Of particular significance, the hypothetical function assigned to pre-frontal activations and their temporal dynamics has played a major role in global theories of consciousness ^12^. For example, “localist” theories (e.g., ^13–15^), posit that different contents of awareness should be linked to information in local cortical regions will argue that the visual pre-frontal regions are likely linked to non-perceptual aspects-typically attributed to the frontal lobe-such as language and decision making. In contrast, “globalist” theories-such as *Global Neuronal Workspace*, or *High Order Theories*^16–19^ will attribute to frontal activations a central role in enabling visual perceptual awareness. Moreover, according to the *Global Neuronal Workspace* theory, prefrontal activations are hypothesized to be transient, showing at both the appearance and disappearance of perceived stimuli, reflecting the updating of consciously perceived contents^8,12^ (for a recent review of this debate see ^20,21^). Thus, highlighting the potential functional role and dynamics of visual activation in human prefrontal cortex, beyond its immediate interest in understanding the functionality of the frontal lobes has also far-reaching theoretical consequences.

In recent years, the striking success of deep neural networks (DNNs) opened a new window for addressing the difficult problem of cortical functionality (see ^22,23^). A particularly fruitful research direction has been the search for parallel relational structures in brains and artificial networks^24–29^ (for review, see ^30^). Relational structures are a proposed representational scheme in which cognitive content, e.g., a visual image, is represented by the set of similarities and differences of this image from other visual images^13,31–33^. Such representations are particularly convenient when comparing different systems and modalities. In the domain of human perception, various recent studies have shown that these algorithms account for a significant proportion of variance in human representations of faces and objects, outperforming any previous computational models^30,34–37^. Given the finding of significant parallels between artificial networks and visual representations at the posterior cortex, it is of interest to search for such parallels in the visually active sites found in the human frontal lobe.

Typically, studies investigating brain activity during perception utilize DNNs that are trained on visual inputs^35,38,39^. In a recent study Shoham et al^40^ proposed a novel approach, involving models trained on different types of information (visual or textual) to test whether they explain unique perceptual and semantic components in the representation of familiar visual stimuli in human perception and memory. In the current study we extended this paradigm to explore brain activity. Specifically, we aimed to discern whether the response of the frontal lobe to visual stimuli is primarily encoding visual or textual information. To achieve this, we employed visual (ImageNet-trained VGG-16) and text (GPT2) DNNs to model brain activity. Recent findings showed that the multi-modal, image-language DNN, CLIP (Contrastive learning image pre-training) also accounts for a significant proportion of variance in behavioral^40^ and neural representations^41–43^. Therefore, in addition to standard visual and language DNNs, we also extracted visual and textual embeddings from CLIP image and text encoders. To quantify the contributions of visual and textual information to visual activations in the frontal lobe, we used intra-cranially recorded iEEG data in which patients were presented with images of famous individuals and places and were asked to memorize the visual details in each image for a later recognition test (see Figure 1). This method enabled us to follow the precise dynamics of these frontal processes during perception and investigate whether they are mainly driven by visual or textual information.

**Figure 1:**
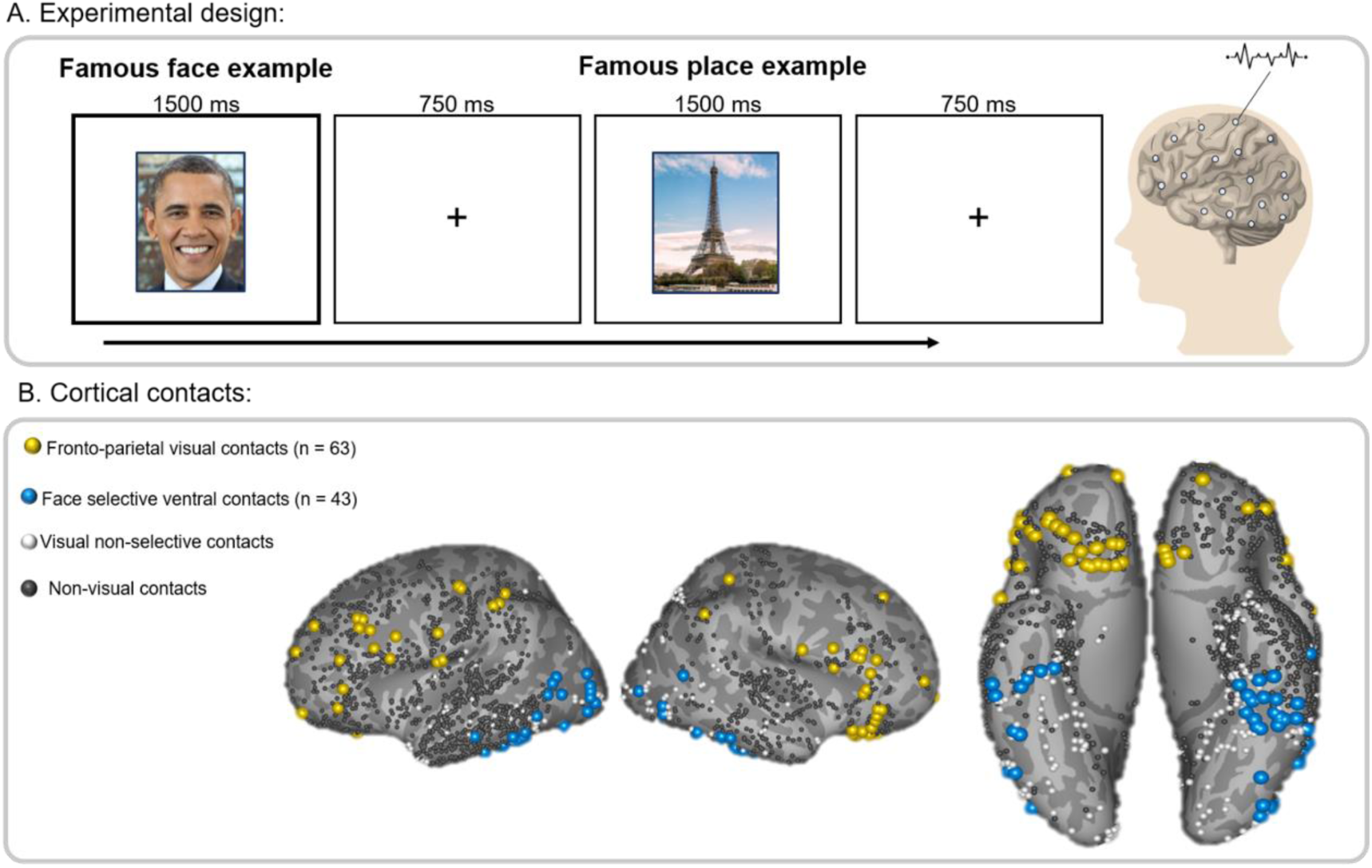
Experimental design and general methods: **a.** Experimental design: participants viewed 28 images of familiar faces or places while their brain activity was recorded via intracranial EEG. **b.** Contact locations: yellow - Fronto-parietal visual contacts, blue – Face-selective ventral contacts, white – visual non-selective contacts, gray- non-visual contacts.

## Results

### Experimental design and intra-cranial contacts information

Our study included intra-cranial recordings, conducted for clinical purposes in 13 patients (10 females, mean age 34.7 ± 9.6) and 2571 recording sites. The participants were presented with images of famous individuals and places. Briefly, participants viewed 28 different images, 14 familiar places and 14 familiar faces, divided to two experimental runs. Each image was presented four times during the run, in a pseudo-random order, ensuring no image was repeated twice consecutively. The images were presented for a sustained time duration of 1500 ms, with 750 ms inter-stimulus fixation intervals. No participants were excluded based on performance (for further details see methods and^44,45^).

Figure 1 depicts the experimental set-up and recording sites that were examined in the present study. We first identified the visually responsive contacts (n=377, out of the 2571 total contacts) and then divided them, based on their anatomical locations (ventral, fronto-parietal) and functional characteristics, into the different regions of interest (ROIs) (see methods of^45^ for details).

### Extraction of visual and textual representations from DNNs

Visual representations of the 28 images were extracted from the final fully connected layer of an ImageNet trained DNN (VGG-16) and from the final layer of the visual encoder of CLIP. Textual representations of the same familiar stimuli were the embeddings of the first paragraph of their Wikipedia description based on the final layer of GPT2, and the embeddings of their names, based on the textual encoder of CLIP (Supplementary Figure 7 shows that CLIP’s name-based representations carry relevant semantic information. See Methods section for details). We then generated RDMs, by computing the cosine distance between the embeddings of the 28 images/textual descriptions for each DNN (see Figure 2A). Resulting in four different RDMs, two visual-based RDMs (VGG, CLIP-Image) and two text-based RDMs (CLIP-Text, GPT). The correlations between the different DNNs are presented in Figure 2B.

**Figure 2:**
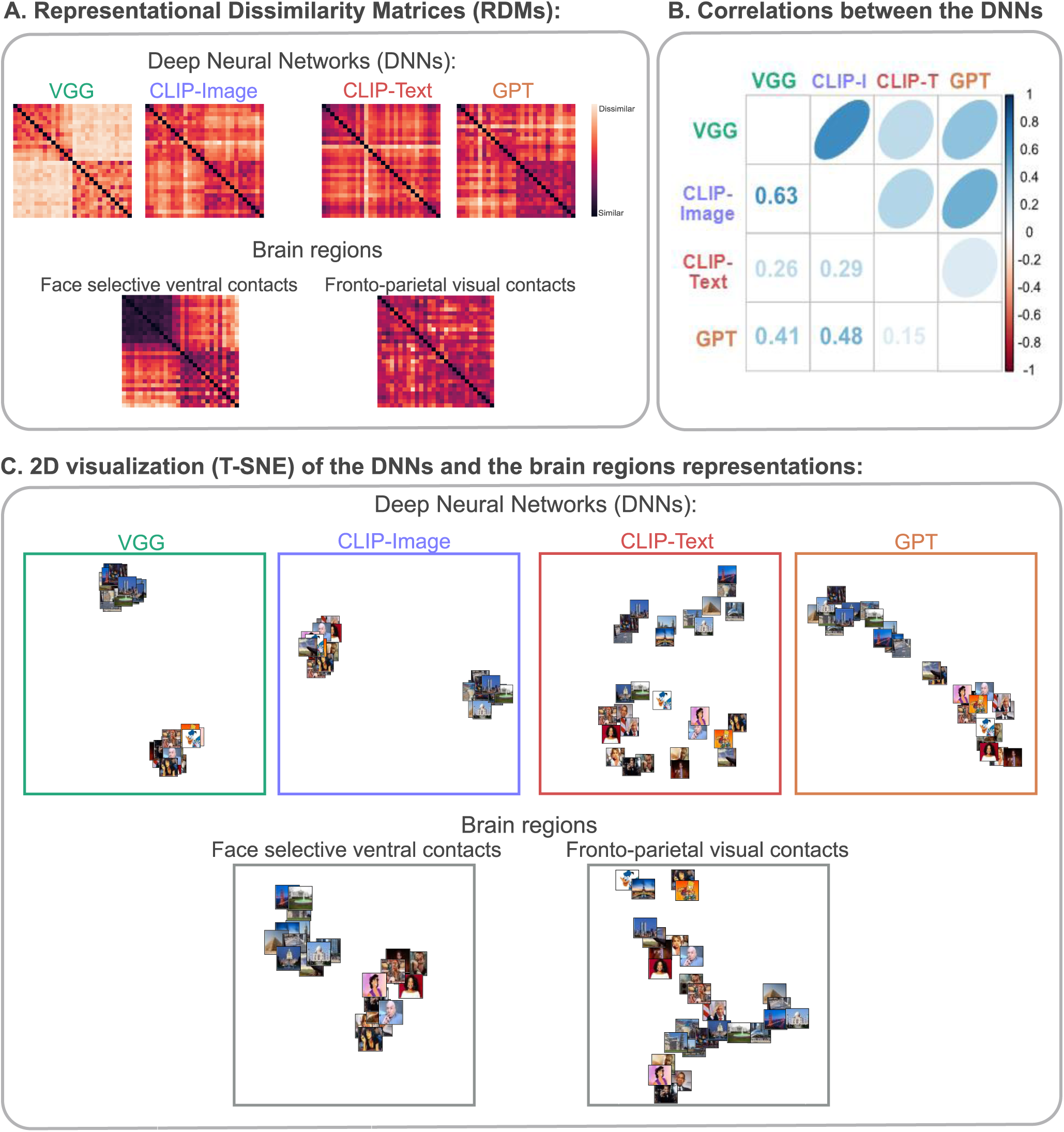
Representational Dissimilarity Matrices and their 2D visualization: **a.** Representational Dissimilarity Matrices (RDMs) of the 28 stimuli. RDMs of the embeddings of the images based on visual DNNs (VGG, CLIP-image) and of their textual description based on textual DNNs (CLIP-text, GPT). RDMs of the neural response to the images in face-selective ventral and fronto-parietal visual contacts. **b.** The correlations between the RDMs of the visual and textual DNNs. **c.** A 2D visualization (T-SNE) of the geometrical representations of the stimuli based on the visual and textual DNNs and the neural response to the images in face-selective ventral and fronto-parietal visual contacts.

### Fronto-parietal visual contacts are correlated with text-based rather than visual-based DNNs

To explore the potential parallels between the visual activations in the frontal lobe and DNNs, we compared their relational structures as revealed through representational dissimilarity matrices (RDMs, see ^34^). For each stimulus, we calculated the average response in each contact within a range of 0.1-0.4 sec (100-400 ms) from the moment the stimulus was presented. We mainly focused on this time window as it encompasses the peak of the response, but we explored the response averaged across other time windows as well (time windows at 0.4-0.7 sec, 0.7-1.0 sec, 1.0-1.3 sec, and 1.5-1.8 sec) (see Fig. 4). We defined the contacts’ mean response (averaged across 4 repetitions) population vectors as the stimulus activation patterns in each ROI and measured the Pearson correlation between each two stimuli activation patterns. One minus correlation represented the dissimilarity score between the two stimuli (lower = more similar) in this ROI. This provided us with the RDM for each ROI (see Figure 2A).

Next, we compared the relational-structure similarities between the fronto-parietal contacts and the output layers of four different deep convolutional networks: two networks were text-based (GPT and CLIP-Text) and two networks were image-based (CLIP-image and VGG). For further details about the different DNNs see methods. The correspondence between the patients’ frontal-parietal ROI RDMs (or the face-selective ventral ROI) and the artificial networks’ layers were computed using Pearson correlations: We computed the correlation between the frontal ROI RDM with each network RDM (Figure 3 top panels). All four networks were correlated with the response of the frontal lobe contacts to the images. However, our aim in this analysis was to examine the unique contribution of each DNN network to the overall variance. To test this unique similarity between each network representation with the frontal lobe we calculated the partial correlations between each DNN and the patients’ RDM, when all other DNNs are held constant. To test for statistical significance, we computed the same correlations using a permutation test (shuffled pairs labels, see methods for further details) and bootstrap analyses (leaving one participant/contact/stimuli/repetition out). We tested both the significance of the predictors’ partial correlations individually and the difference between each partial correlation in each ROI. P-values were FDR corrected for each ROI and each analysis (individual predictors and differences) separately.

**Figure 3:**
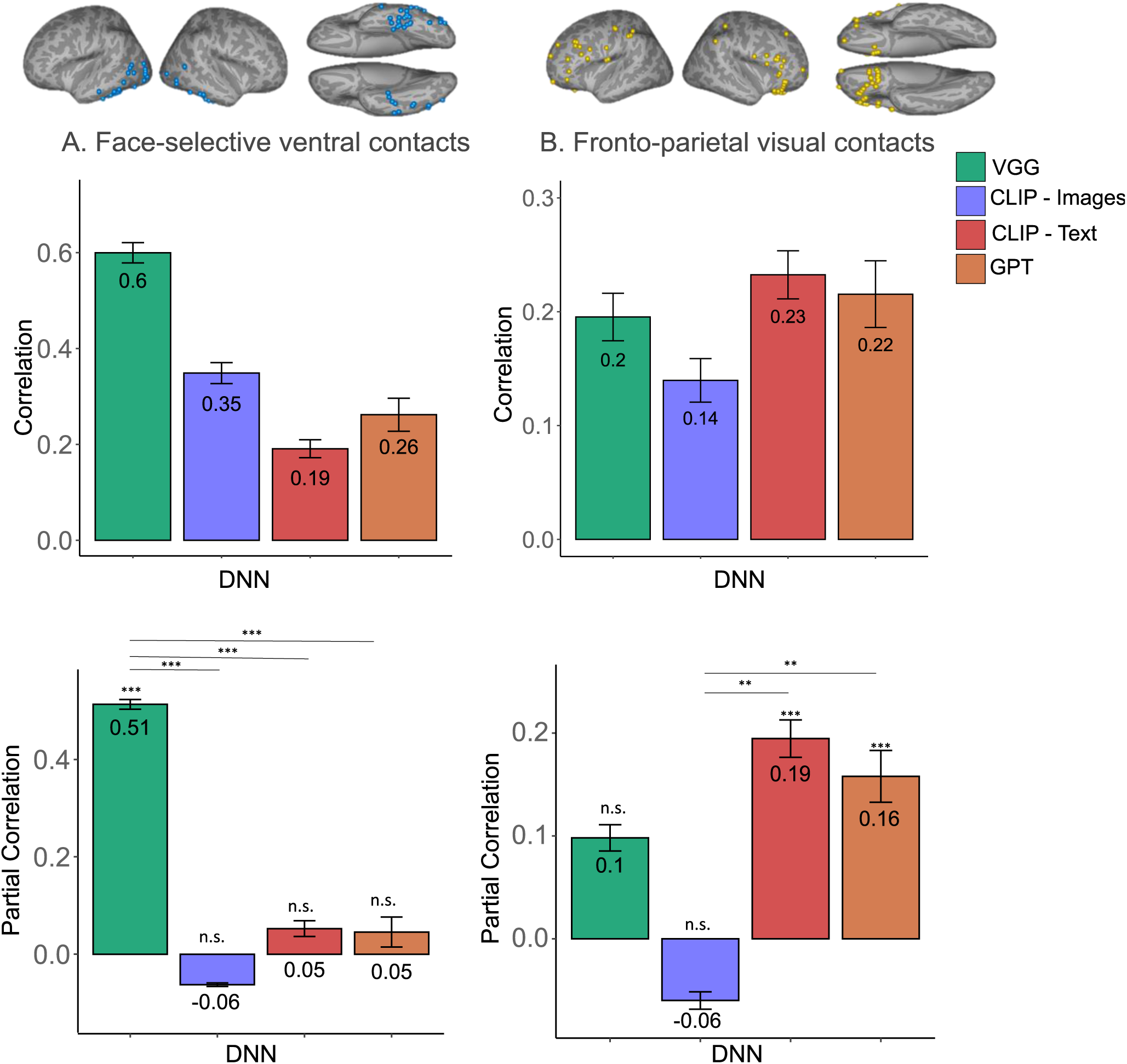
Correlations (Top) and Partial correlations (bottom) of image and text DNNs with Face-selective ventral and Frontal-parietal visual contacts: Zero order correlations are presented in top panels and partial correlations between each of the DNNs, when the other three DNNs are held constant, are presented in bottom panels. **A**. Face-selective occipito-temporal contacts (n=43), Error bars indicate leave one participant out procedure s.e.m. **B.** Fronto-parietal visual contacts (n=63), Error bars indicate leave one participant out procedure s.e.m. All *p* values were derived from a pair-images permutation test (10000 permutations). Reported *p* values are FDR corrected in each ROI separately. Note the clear shift in bias from visual-based to text-based preference from occipito-temporal to frontal-parietal ROIs. ∗∗ 𝑝_𝐹𝐷𝑅_ < 0.01, ∗∗∗ 𝑝_𝐹𝐷𝑅_ < 0.001

Figure 3 (bottom panels) depicts the partial correlations found between each of the different DNNs (warm colors depict the text-related while cold colors the visual-related ones), when the other three DNNs are held constant. The figure shows the occipito-temporal, high order, visual face-related ventral ROI (Panel A bottom) and the fronto parietal visual contacts (Panel B bottom) in the 0.1-0.4 sec time window. As can be seen, in the Fronto-Parietal ROI, only the text-related DNNs displayed significant partial correlations to the patients’ RDMs. The opposite selectivity was found in the face-selective ventral contacts in the visual cortex. Here, only the visual-related VGG DNN showed a significant correlation to the Face-selective ventral contacts.

As shown in Figure 3B top panel, all four networks were correlated with the Fronto-Parietal ROI. To better understand whether the visual networks (whose correlation dropped in the partial correlation analysis), account for the same information as the text networks, we added a category RDM as a fifth predictor in the partial correlation analysis. The category RDM was implemented by assigning similarity scores of 0 for same-category pairs (face-face or place-place) and 1 for different-category pairs (face-place). We then calculated the partial correlations between each predictor and the Fronto-Parietal ROI. When adding the category RDM as a predictor, the visual networks showed very low and non-significant partial correlations. In contrast, the category predictor did not alter the significant association between the Fronto-Parietal ROI and the text-based DNNs (see Supplementary Figure 1). These findings suggest that the association of the text-based networks with the Fronto-Parietal ROI is not category-based.

Finally, given that the stimuli set in this study included both places and faces, we employed a second control ROI that was not only face-selective but rather included contacts in high-order visual cortex that showed significant preference for a subset of the stimuli (could be faces, places, or a mix of both-Content Selective ROI, see Supplementary Material). The pattern observed in this ROI was similar to the results found for faces-selective ventral contacts (for details, see Supplementary Material and Supplementary Figure 2).

### Sustained textual contribution with a stimulus-offset response in frontal-parietal visual responses

Given that CLIP-Text showed the strongest partial correlation with the fronto-parietal visual contacts, we proceeded to investigate this brain-network correlation across different time windows (Figure 4). For comparative analysis, we examined CLIP-Text correlations within the same time windows with face-selective ventral contacts as well as the correlations of the ventral and frontal-parietal contacts with the visual DNN (VGG) (Note that analysis shown in Figure 4 displays the correlations and not the partial correlations that are shown in Figure 3 bottom panels). To assess the significance of differences in correlations across time windows, we used a permutation test (see methods for further details). When exploring CLIP-Text and fronto-parietal correlations (Figure 4, bottom-right panel), they were all as high as the correlation of the first time window (0.1-0.4 sec) besides the 1.0-1.3 sec, which showed no correlation. This might suggest a relatively persistent processing pattern during the stimuli presentation until 1 sec.

**Figure 4:**
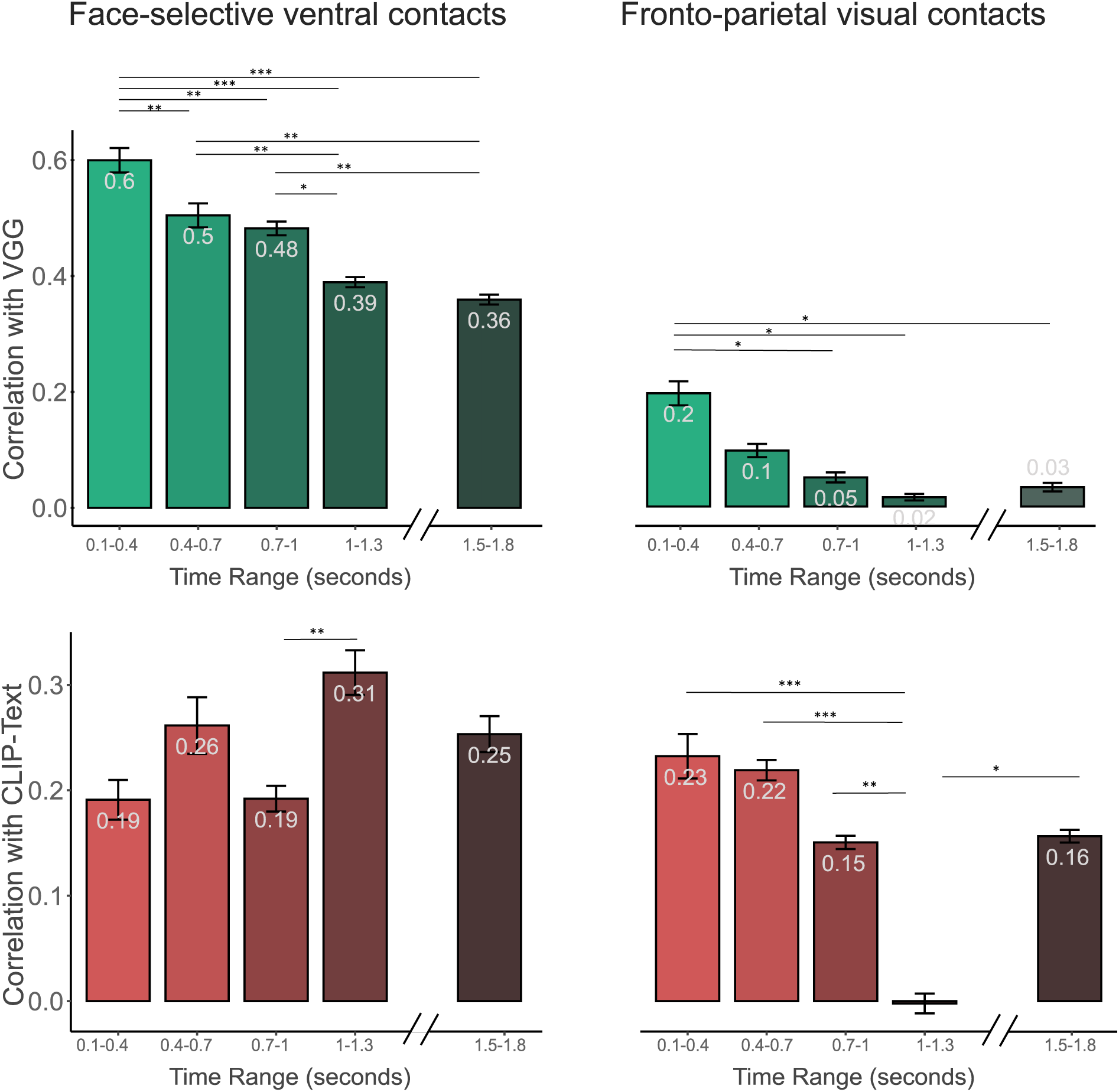
Correlations between VGG or CLIP-Text with Face selective ventral and Frontal-parietal visual contacts at different time windows: Top bar plots (green bars) show the correlations with VGG in face-selective ventral contacts (left), and fronto-parietal visual contacts (right). Bottom bar plots (red bars) show the correlations with CLIP-Text in face-selective ventral contacts (left), and fronto-parietal visual contacts (right). Note the strikingly different dynamics in fronto-parietal contacts between the visual component that declined rapidly and the text component with persisted followed by an offset response. Error bars indicate leave one participant out procedure s.e.m ∗ 𝑝_𝐹𝐷𝑅_ < 0.05, ∗∗ 𝑝_𝐹𝐷𝑅_ < 0.01, ∗∗∗ 𝑝_𝐹𝐷𝑅_ < 0.001

Interestingly, a similar pattern seems to reappear at the stimulus offset (1.5 sec). This was not the case for VGG correlations with the fronto-parietal visual contacts (Figure 4 top-right panel), which showed a steep decline after the first-time window. Nor is the case for the face-selective ventral contacts that showed gradual decline in VGG correlations (Figure 4 top-left panel) and variable CLIP-Text correlations with time (Figure 4 bottom-left panel). see Supplementary Figure 3 for a similar analysis with GPT.

### A monotonic increase in the correlations with fronto-parietal visual contacts across the layers of CLIP-Text

To further explore the initial-time window (0.1-0.4 sec) we asked whether the frontal correlation is merely due to the inner structure of the DNNs, regardless of its input? We also asked how it is manifested across the different layers of the DNN. Figure 5 shows that the correlation to CLIP-Text based on shuffled text input is drastically reduced compared to the original text (the ‘output’ layer was used in the previous analyses; see Supplementary Figure 8 for the same comparison between GPT’s embeddings based on the original and shuffled text, which shows similar results). We also extracted the RDM according to each layer of CLIP-Text network (see methods for further details). Examining the evolution of the correlation to the Frontal contacts’ RDM through the DNN layers revealed a clear monotonic increase towards higher order layers, reaching the highest values at the three top layers (Residual attention blocks (RAB) 11,12, and output-13), indicating a specific, functional similarity between the frontal lobe responses and the top CLIP-Text layers. Also note the striking difference between the RDM correlations of the correct and shuffled texts.

**Figure 5:**
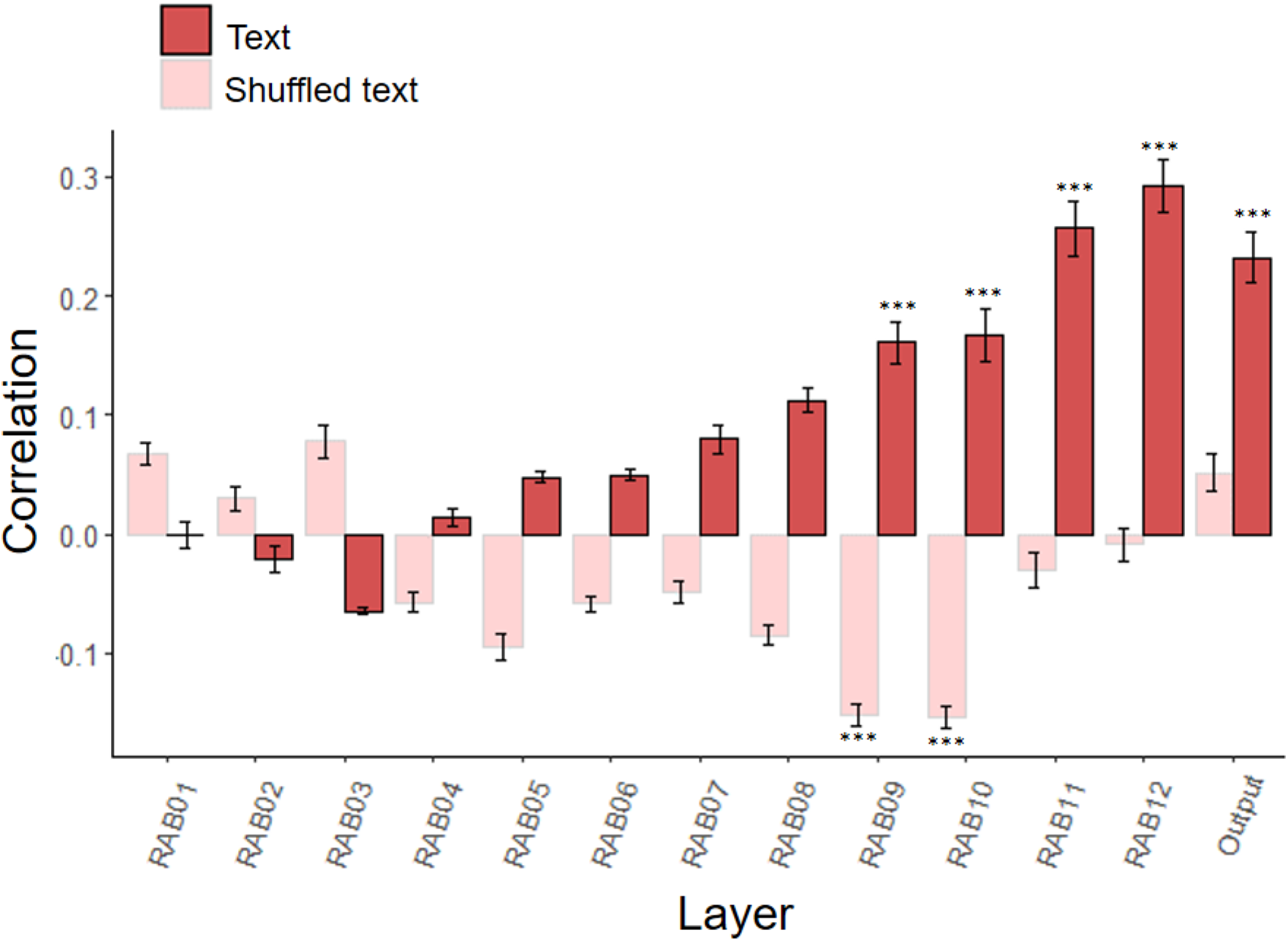
CLIP-Text different layers’ correlations with Fronto-parietal visual contacts when embeddings were based on the original text or on shuffled text: The correlations with the fronto-parietal visual contacts in the first time window (0.1-0.4 sec) with CLIP-Text different layers based on the original text (opaque) or shuffled text (pale). Error bars indicate leave one participant out procedure s.e.m. ∗∗ 𝑝_𝐹𝐷𝑅_ < 0.01 ∗∗∗ 𝑝_𝐹𝐷𝑅_ < 0.001

To rule out the possibility that these correlations were a result of an outlier patient, contact or stimulus response, we examined how robust was the correlation to CLIP-Text when using only partial data in the analysis. These analyses and results are reported in the Supplementary material. As can be seen in Supplementary Figure 4, the results remained fairly robust to different bootstrap analyses, arguing against an outlier-driven effect.

### Correlations of CLIP-Text with fronto-parietal visual contacts are maximal for the first image repetition

Since in the experiment the same visual images were shown four times to each patient, it was of interest to examine how such repetitions affected the observed correlation to the CLIP-Text network. To that end, separate RDMs were constructed for each repetition of the stimuli. Supplementary Figure 5 depicts the RDM correlations across CLIP-Text layers for each of the repetitions. Result show that the correlation levels changed in a complex manner across repetitions. With the novel images (first appearance) failing to show any correlation and a significant correlation appearing for the first repetition followed by a progressive decline in the second and third ones.

Finally, to examine whether the correlation to CLIP-Text changes across different frontal subdivisions, the contacts were divided into two major anatomical subdivisions, orbito-frontal and lateral-frontal contacts (see further detailed in the Supplementary file). As can be seen in Supplementary Figure 6, the overall pattern of correlations across anatomical regions in the frontal cortex remained largely unchanged showing a gradual increase in correlation when moving from low-level to high-level network layers.

## Discussion

In this study, we used text-based and image-based DNNs to investigate their similarity to brain activity, while participants viewed images of famous faces and places. Our analysis revealed that visually responsive fronto-parietal contacts’ activations were best predicted by text-based networks, suggesting semantic rather than visual-related processing.

### The frontal lobe processing in visual perception

Our results reveal, in agreement with a number of prior studies based on both fMRI and intracranial brain recordings^2,3,8–11,46^, that the frontal lobes are consistently activated by visual images. Importantly, in the present study the patients were instructed to view and memorize the images rather than report on seeing them or not - so the visual activations cannot be attributed to active reporting but to the perceptual processing of these images.

Though previous studies showed activation in the frontal lobe while viewing visual stimuli^3,8–10,47^, the necessity of the frontal lobe in conscious perception is still unclear. Some research highlights the non-visual nature of frontal lobe tasks, with the highest activations during reporting and motor tasks^2,5,20,48^, and lesion cases challenge the idea of the prefrontal cortex’s essential role in perception^49,50^. It is important to note that identifying the function of cortical responses is a fundamental challenge in all human brain studies, particularly since causal manipulations are rare and necessarily associated with poorly controlled clinical cases. We and others have recently proposed^22,40,51,52^ that insight into the functionality of cortical responses can be gained by searching for “convergent evolution”- i.e. parallels that are found between artificial networks and brain systems^23,34,53^. Here we adopted this strategy to gain insights into the enigma of frontal visual responses. By searching for correlations between relational structures, revealed through RDMs, in frontal cortex and visual and textual deep learning networks, we have uncovered a preference for text-related networks. The correlations to the image-based networks although positive, were far weaker and did not account significantly for the visual frontal response beyond the textual-based networks.

The fact that the more specific correlation was found to a text-based network rather than a similar visually related ones (CLIP-image) is intriguing, especially considering that the patients’ RDMs were based on visual responses and that visual-related DNNs were correlated with visual contacts in high level visual cortex. This finding suggests that the frontal visual responses may be more closely tied to lingual analysis influenced by visual contents rather than visual perception per-se. Notably, robust visual responses associated with non-perceptual functions are well-documented in the human brain. Particularly striking cases are visual activations found in the hippocampus associated with conceptual representations^54,55^ and visual activations in human high order somatosensory cortex^56,57^. Given the established role of the frontal lobes in language and thought processing^58^, it is not surprising that the visual activations in this study appear to be more linked to linguistic rather than perceptual processing functions.

### The temporal dynamics of prefrontal and posterior correlations

Comparing the temporal dynamics of correlations between the RDMs of DNNs and brain activations revealed different patterns of results in posterior compared to frontal ROIs. In the posterior face selective ventral ROI both image-based (VGG) and text-based (CLIP-Text) networks showed consistent correlations across time windows, with higher correlations for VGG. Conversely, activations in the frontal ROI were correlated with VGG only in the earliest time window (0.1-0.4 sec), in sharp contrast to the correlations profile of CLIP-Text which were found to be significant in all but one of the time windows (1-1.3 sec). These results echo previous findings showing persistent representations of visual contents in posterior regions^8–10^, and add novel insight regarding the dynamics of prefrontal activations in two ways: First, the obtained results point to a differential dynamics for the transient, visual-related aspects of the frontal activations vs the persistent text-related ones. Second, by focusing on the Text-based component of the frontal activation, our results demonstrate an intriguing offset response, predicted, albeit for visual processing (up to 300 ms after stimulus offset), by the Global Neuronal Workspace theory^8^. It is worth mentioning that whereas both CLIP-Text and GPT were similarly correlated with frontal-parietal contacts during the early latency, they exhibit distinct patterns of correlation when looking at later time windows (Figure 4 and Supplementary Figure 3). These variations might be due to their different input (name in Clip-Text or paragraph from Wikipedia in GPT) or the way they were trained. CLIP-Text is a multi-modal visual-semantic DNN, while GPT is a unimodal language DNN. Further research is needed to account for these differences.

### Dynamic modulation of correlations by stimulus repetition

The experimental design in the present study included four repetitions of each of the stimuli presented. The temporal distance between repetitions was randomized across stimuli-however, grouping the repetitions according to their sequential repetition order revealed a complex dynamic of the correlation magnitude to the CLIP-Text network. Thus, one could discern two, not necessarily linked, changes (see Supplementary Figure 5). The first was a rapid increase from essentially no correlation in the first, novel image presentation to the first time the images were repeated. This suggests that the frontal cortex function that is correlated to the CLIP-Text network may not be an encoding function, but rather related to short-term memory or recall. It is interesting to note that a similar discrepancy between novel and repeated presentations were observed in the hippocampal formation^44^ where content-selectivity appeared only on a second presentation, while the first response was a more generalized novelty signal. It is intriguing to speculate that this novelty effect may have been dominant also in the pre-frontal recordings and masked the potential correlation with the CLIP-Text network. The second effect was a gradual decline in the correlation at further repetitions. This type of gradual decline, termed adaptation or repetitions suppression, has been ubiquitously observed in the visual system^9,59–62^. It is interesting to note that what appears as a repetition-suppression effect can be revealed through the window of brain-artificial network correlations.

### Sources for the low-level correlation magnitude

Examining the magnitude of the correlation between the RDMs derived from frontal activations and the CLIP-Text layers even when specifically focusing on the optimal conditions, failed to reveal correlation levels higher than 0.3. This is a significantly lower value compared to magnitude of the brain to network correlation values found in high order visual areas – which was substantially higher, ranging from 0.4-0.6^34^ (and see VGG correlations in Figure 4). Therefore, it could be argued that the differential correlation pattern we found in frontal and posterior areas merely reflects a coarser grained representation in frontal cortex. However, the finding that text-models show higher correlation in frontal compared to posterior visual areas on the one hand, and the discovery of different temporal dynamics of the correlations in frontal vs posterior regions-argue against such possibility. Moreover, while the frontal correlation of 0.2-0.3 was lower than the visual correlation (0.6) – it should be noted that such correlations are consistent with a number of previous studies showing similarly lower, yet significant, correlations ^63–70^.

A number of sources, not necessarily mutually exclusive, could account for such relatively low correlation values. The most obvious factor is the relatively low activation level of the frontal contacts. This weak activation and low SNR is typical in report-free paradigms such as the one adopted in the present study (see also ^2,9^). Whether this effect is simply due to smaller signal to noise ratios in the weakly responding contacts or whether it reflects a less selective signal in these contacts remains to be studied.

A second possibility is that as indeed suggested by the preferential correlation to the text-related network the true function of the pre-frontal contacts is not perceptual but language related. Since the task of the patients did not involve explicit language or textual aspects-the visual activation may have merely occurred as an automatic, auxiliary by-product of the visual presentation and memorization of a familiar image - a ubiquitous cortical phenomena as has been pointed out above^20,71,72^. Resolving this issue will necessitate additional experiments in which linguistic and decision-making tasks will be manipulated in addition to the visual images.

A second limitation to note is that we used a relatively small number of stimuli (n = 28), whereas recent fMRI studies have explored neural representations with a much larger set of stimuli^41,42^. iEEG is a superior method compared to non-invasive techniques such as fMRI or scalp EEG, as it combines high temporal resolution, allowing us to explore, dynamic changes in correlations, with high spatial resolution, which is more directly related to underlying neuronal activity. However, a major limitation of this unique method is that due to patient fatigue an extensive study including many images and categories is not feasible. Previous studies explored brain representation using different recording methods, have also used such limited set size of stimuli^34,63–65,69,70,73^.

A number of recent studies^41,42,74–76^ have reported that the representations of textual descriptions of visual images predict the visual responses to these images in visual cortex. Note that the textual descriptions used in our study are not descriptions of the images visual appearance but describe their conceptual meaning (names/Wikipedia definition). In an ongoing study, we are currently exploring the possibility that the type of textual material used in describing the visual images may be manifested differently in ventral and frontal brain activations

Finally, one cannot rule out the possibility that more advanced network structures may produce higher and more consistent activation levels. The recent explosive growth in the variety and power of language models will offer rich ground for future explorations in this domain.

To conclude, our findings suggest a closer association between visually responsive fronto-parietal regions and text-based rather than visually related DNNs. The temporal dynamics of this text-based association was observed in different latencies of the stimuli presentations with a drop and a recovery after the stimulus offset, which is aligned with the Global Neuronal Workspace theory. However, our findings highlight the non-visual processing of the frontal lobe during perception, arguing against global theories of consciousness which posit that visual processing should spread to fronto-parietal cortex during conscious vision.

## Methods

### Human data

We describe here the participants tasks and stimuli as well as ROI definitions. Additional details can be found in^9^.

#### Participants

Electrophysiological data were acquired from 13 patients (10 females, mean age 34.7 ± 9.6) via intracranial iEEG recordings. The recordings were obtained while the patients underwent pre-surgical evaluation for drug-resistant epilepsy at North Shore University Hospital in NY. As part of the clinical assessment, subdural or depth contacts were implanted in all patients. The study followed the latest version of Declaration of Helsinki, and all patients provided a fully informed consent to participate, including consent to publish the results, according to the US National Institute of Health guidelines. The Feinstein Institute for Medical Research’s institutional review board monitored the study. Notably, no clinical seizures occurred during the experiment.

#### Experimental task and stimuli

The experiment comprised two runs, each starting with a 200-second resting phase with closed eyes. Subsequently, participants viewed 14 different images in each session (7 images of each category – famous faces or famous places). In total, 28 stimuli were used, 14 in each run (a similar stimuli set size, was used previously in studies that measured neural representations of visual stimuli^34,63–65,69,70,73^). Each image was presented four times (for 1500 ms each time, with 750 ms inter-stimulus intervals) in a semi-random order to avoid consecutive repeats. Participants were informed that they would later be required to recall the images and describe their visual features in detail, rather than merely identifying them. Consequently, they were instructed to memorize the details of the images. Participants performed well, and no participants were excluded based on performance (Further details can be found in ^45^).

#### Definition and grouping of visually responsive contacts

Our study comprised of 2571 recording sites (contacts). However, the analyses focused only on visually responsive contacts, subdivided into different ROIs. Visual responsivity was defined as electrodes displaying statistically significant activations in response to stimuli presentations. To do so, the data from all contacts (visual and non-visual) was preprocessed, and transformed to the high-frequency broadband (HFB) signal (see ^9,44,45^ for further details). We extracted visual-responsive contacts by a comparison of each contact’s post-stimulus HFB response, averaged across the 100 to 500 ms period following stimulus onset, with its pre-stimulus baseline, averaged across the −400ms to −100ms interval prior to stimulus onset. This comparison was conducted using a two-tailed Wilcoxon signed-rank test, that was FDR corrected for all contacts (from all patients) together. Subsequently, contacts demonstrating a significant HFB response (𝑝_𝑓𝑑𝑟_ < 0.05) were categorized as visually responsive, totaling 377 contacts.

Following this, visually responsive contacts were grouped into subsets based on their anatomical and functional characteristics, namely face-selective ventral and fronto-parietal visual contacts. Face-selective ventral contacts were defined by comparing mean HFB responses between faces and places during the 100-500ms post-stimulus window using a Wilcoxon rank-sum test. Contacts showing significantly greater activation to faces compared to places (𝑝_𝑓𝑑𝑟_ < 0.05), located anatomically beyond early visual regions (but excluding the frontal cortex), were categorized as face-selective ventral contacts (n = 43, depicted in blue in Figure 1B). We defined the fronto-parietal ROI based on the Desikan Killiany atlas locations of the following labels: superior frontal gyrus, rostral middle frontal gyrus, pars orbitalis, pars triangularis, pars opercularis, precentral gyrus, postcentral gyrus, supramarginal gyrus, orbital frontal gyrus, and the anterior cingulate. Visual contacts located within these regions were categorized as the fronto-parietal visual contacts (n = 63, depicted in yellow in Figure 1B). Any visually responsive contacts not falling into these specified categories were termed “other visually-responsive contacts” and are depicted in white in Figure 1B, while non-visual contacts are depicted in gray. Further details can be found in ^9^. A third group of visual contacts was extracted, to be used as a second control for the Fronto-parietal ROI. We refer to this group as content selective contacts (n=114). Their extraction and results are fully described in the Supplementary Material and Supplementary Figure 2.

#### Deep neural networks

Visual network – ImageNet pre-trained VGG-16: To examine a network based visual representation of the presented stimuli, we used the VGG-16 network^77^ pre-trained on ImageNet^78^, which includes 300000 images from 1000 object categories. VGG is a commonly used network performing object and face categorization at human level^40,79–81^. We then extracted the embeddings of each image based on the feature vector representation in the penultimate- i.e. the hidden layer just below the output (fc7) layer of the network, which is the representation that is used for the classification. The similarity between each images pair was computed based on the cosine distance between these feature vectors.

Text network – openAI GPT2: A transformer-based language algorithm^82^. GPT is trained on a vast corpus of text to carry out various language tasks (e.g. semantic similarity, questions answering, grammatical correction etc.), often involving longer text captions. Therefore, we chose to extract its representations using text from Wikipedia to obtain the representation of each stimulus (see Supplementary Table 1 for the text used for each stimulus). Specifically, we retrieved the embeddings of the first paragraph from Wikipedia corresponding to each familiar stimuli (identity or a place), based on the model output layer. Subsequently, we computed the similarity between each stimuli pair based on the cosine distance between the representations of these textual descriptions.

Multi-modal network - CLIP (Contrastive Language-Image pre-training)^83^: CLIP is trained in a self-supervised, contrastive manner to create similar representations for images and their text caption based on a training set of 400M images and their captions from the internet^83^. We extracted the embeddings of each image based on the output layer of trained ResNet architecture using the visual and language components of CLIP.

CLIP Visual Component (CLIP-Image): We extracted the representations of the 28 visual images that were presented to the patients. We then computed the similarity between each object pair based on the cosine distance between these visual embeddings. We will refer to these representations as CLIP-Image.

CLIP Text component (CLIP-Text): CLIP textual component is trained with short captions that are used as labels of images on the internet and is limited to short text input (up to 77 characters). Therefore, we extracted its representations based on the names of the stimuli (see Supplementary Table 1 for the name used for each stimulus). We extracted the representations based on the names of the familiar exemplar (identity or a place) presented in each stimulus (e.g. “Eiffel Tower”). We computed the similarity between each object pair based on the cosine distance between these name representations. We will refer to these representations as CLIP-Text.

This provided us with the RDM of each network that is based on its output (or penultimate) layer. In addition, we also extracted the RDMs of each layer of the network.

As described above, we extracted the representations of GPT and CLIP-Text based on different text inputs. The representations of GPT were based on Wikipedia definitions, while the representations of CLIP-Text were based on the image’s names. While we chose each input based on the network’s nature of leaning, extracting the representations of CLIP-Text based on names, might seem like a significant reduction in information compared with GPT’s input. However, we believe this is not the case. First, we extracted the similarity scores between all images and all names of the identities/places presented in the images using Image and CLIP-Text, respectively. This enabled us to create a familiarity RDM and identify which name was most similar to each image based on CLIP representation (see Supplementary Figure 7). We found that for each image, the most similar name was its corresponding name (e.g., Barack Obama’s image was closest to the text “Barack Obama” compared to any other name in the list), suggesting that CLIP’s name-based representations carry relevant semantic information. Moreover, as seen in the results section, these CLIP-Text representations had the highest association with the frontal contacts, indicating that they represent the stimuli in some informative way. Finally, to test if we could extract further information from CLIP-Text, we also extracted its representations based on the Wikipedia definition (limited to 77 characters), which did not improve its association with the brain ROIs (see Supplementary Material for details).

### Data Analysis

For the purpose of Representational Similarity Analysis (RSA) we created a dissimilarity matrix (RDM) of all possible stimuli pairs within the pool of the 28 stimuli (a total of 378 pairs), separately for each different brain ROI and the different DNNs.

We first computed the Pearson correlation between the RDMs of each DNN and the RDMs of the ventral and frontal visual contacts. Then, to assess the unique contribution of each DNN to the neural responses we computed the partial correlation of each DNN while holding the other DNNs constant. Applying partial correlations provide us with the unique association between each pair of variables, that is not mediated by other variables, and this process is not affected by the order of the variables ^84^.

To test the significance of a correlation/partial correlation of a single network in a specific ROI we used a permutation test in which the networks’ similarity scores (between pair of stimuli) were held constant, but we shuffled the similarity scores that are brain based. For each analysis and each ROI, we shuffled the similarity between pairs of stimuli in 10000 iterations. Then we calculated the correlation/partial correlation (respectively) of the network with the patients’ RDM (based on the shuffled similarities). P-value assigned to each network and layer was the proportion of iterations in which the original correlations/partial correlation (based on the real similarities) was greater (or smaller) than the value that was calculated in each iteration. P values were adjusted according to Phipson and Smyth^85^ correction for permutation tests, and then they were FDR corrected for all networks that were tested in the same analysis, for each ROI separately. Then we used a two-tailed test to infer significance. We also tested the significance of the difference between each pair of two DNNs or two time windows; To do so we used a permutation test in which the networks’ similarity scores were held constant, but we shuffled the similarity scores that are brain based. For each analysis and each ROI, we shuffled the similarity between pairs of stimuli in 10000 iterations. Then we calculated the correlation/partial correlation (respectively) of each network/time window with the neural RDM (based on the shuffled similarities), and proceeded to calculate the difference between each two networks/time windows correlations/partial correlations (with the neural RDMs). P value assigned to each comparison of a pair of networks was the proportion of iterations in which the original difference in correlations/partial correlation (based on the real similarities) was greater (or smaller) than the value that was calculated in each iteration. P values were adjusted according to^85^ correction for permutation tests, and then they were FDR corrected for all networks that were tested in the same analysis, for each ROI separately. Then we used a two-tailed test to infer significance.

## Acknowledgments

The study was funded by a CIFAR BMC fellowship to R. Malach and ISF grant no.917/21 to G. Yovel

## Author contributions

Conceptualization – A.S., R.B.D, I.Y, R.M, G.Y; Methodology A.S., R.M, G.Y; Software – A.S., R.B.D; Formal Analysis A.S., R.B.D; Data Curation – A.S. R.B.D; Writing – Original Draft – A.S., R.M.; Writing – Review & Editing – A.S., R.B.D, I.Y, R.M, G.Y; Visualization – A.S.; Supervision - R.M, G.Y; Project Administration - - R.M, G.Y; Funding Acquisition R.M., G.Y

## Supplementary material

Supplementary Figure 1 shows the partial correlation analysis in the Fronto-Parietal ROI when using the RDMs of all 4 DNNs (see main text results section and Figure 3 for the partial correlations including only the 4 DNNs) and a category RDM. The category similarity scores were 0 for same-category pairs (face-face or place-place) and 1 for different-category pairs (face-place). In this analysis, we calculated the partial correlations between the RDMs of each predictor and the RDMs of the Fronto-Parietal ROI. As can be seen, when adding the category RDM as a predictor, the visual DNNs showed very low and non-significant partial correlations with the Fronto-Parietal ROI. In contrast, the text DNNs remained significant, suggesting that the association of the text-based networks with the Fronto-Parietal ROI is not category-based.

**Supplementary Figure 1:**
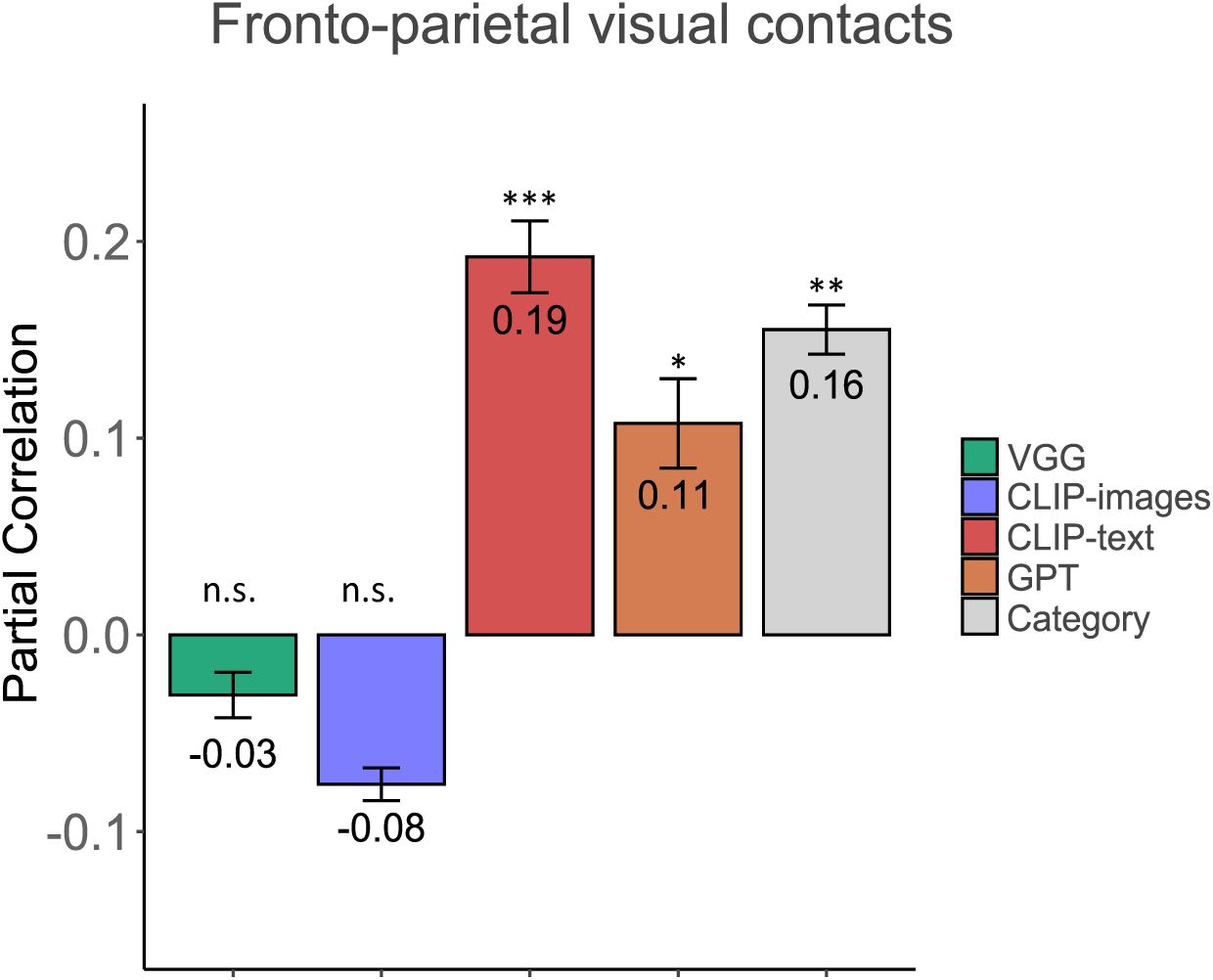
Adding the category RDM as a predictor did not alter the significant association between text-based DNNs and the Fronto-parietal ROI: Partial correlations of the RDMs of image and text DNNs with Frontal-parietal visual contacts when adding the category RDM as fifth predictor: Error bars indicate leave one participant out procedure s.e.m. All p values were derived from a pair-images permutation test (10000 permutations). Reported p values are FDR corrected. The partial correlations of the text-based DNNs with the Fronto-parietal ROI remained significant after adding the category. ∗ 𝑝_𝐹𝐷𝑅_ < 0.05,∗∗ 𝑝_𝐹𝐷𝑅_ < 0.01, ∗∗∗ 𝑝_𝐹𝐷𝑅_ < 0.001

After extracting the visual responsive contacts (see main text Methods section) an additional set of visual contacts was used as a control for the Fronto-parietal ROI, termed content-selective ventral contacts. To define these contacts, we examined the visually responsive contacts in six anatomical regions across the ventral visual hierarchy, based on the Desikan Killiany atlas. These regions included the lateral occipital cortex (LO), inferior temporal gyrus (ITG), lingual gyrus, parahippocampal gyrus (PHG), fusiform gyrus, and entorhinal gyrus, excluding early-visual contacts. Contacts in these regions that showed significant content selectivity (defined as a difference of at least 3.5 SD between the top 10 preferred images and bottom 10 images), were included in the content-selective contact set (n = 114). Twenty-seven contacts met the criteria for both the face-selective and content-selective groups.

**Supplementary Figure 2:**
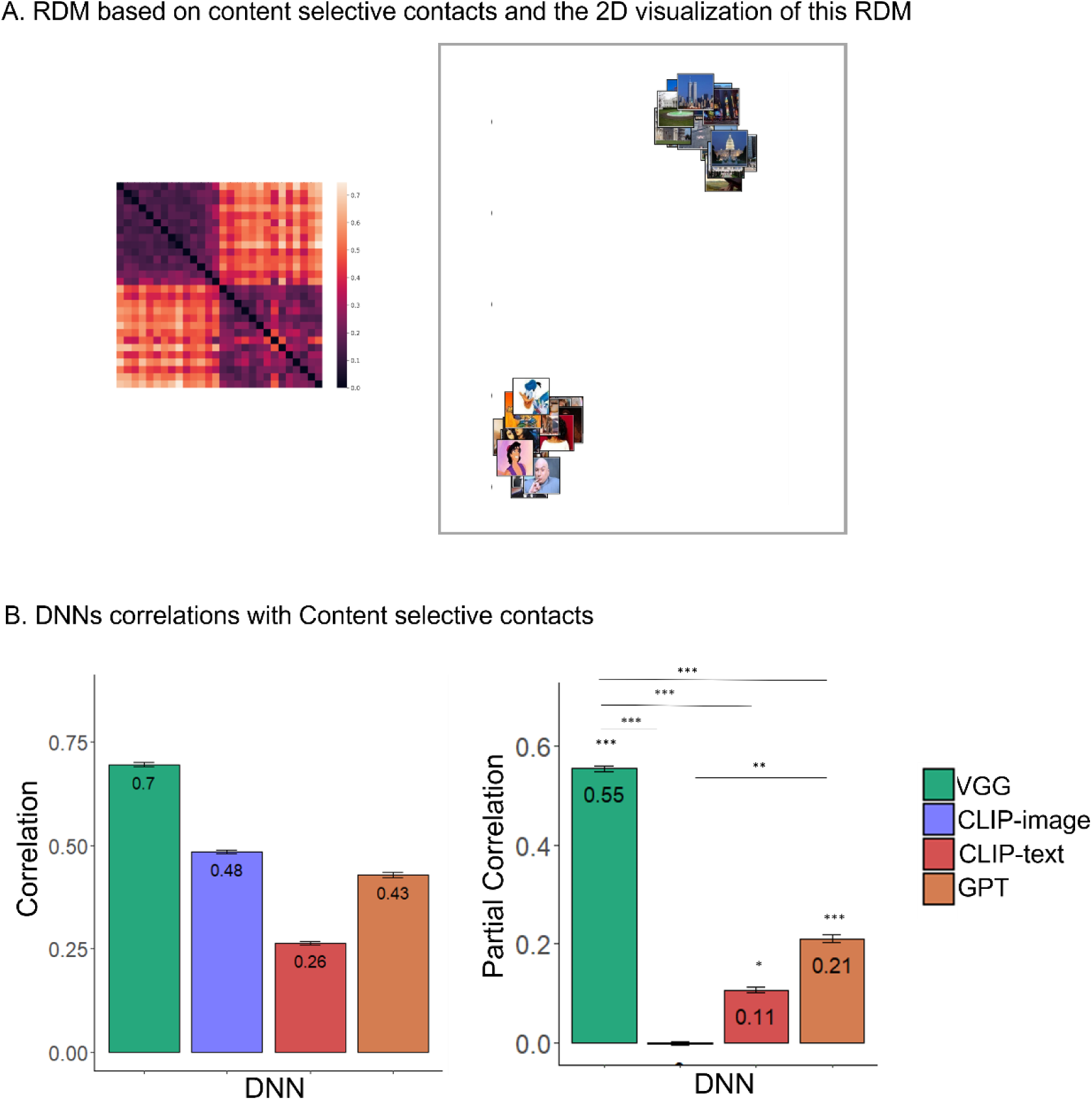
Content selective ROI results: **a.** Representational Dissimilarity Matrices and 2D visualizations: Representational Dissimilarity Matrices (RDMs) that were used for relational structure analysis. **b.** Correlations and Partial correlations of image and text DNNs with a second control for the Fronto-parietal ROI: Zero order correlations are presented on the left and partial correlations between each of the DNNs, when the other three DNNs are held constant, are presented on the right. Error bars indicate leave one participant out procedure s.e.m. The pattern is similar to what we found for Face selective contacts. All *p* values were derived from a pair-images permutation test (10000 permutations). Reported *p* values are FDR corrected in each panel separately. ∗ 𝑝_𝐹𝐷𝑅_ < 0.05 ∗∗ 𝑝_𝐹𝐷𝑅_ < 0.01, ∗∗∗ 𝑝_𝐹𝐷𝑅_ < 0.001

Supplementary Figure 2A shows the RDM that is based on content-selective ventral contacts (the distances were calculated similar to the contacts reported in the main text, see Methods section), and its 2D visualization. Supplementary Figure 2B shows the correlations between the ventral visual content selective ROI and the different DNNs. The left figure shows the zero order correlations, and the right figure shows the partial correlations found between each of the DNNs (warm colors depict the text-related DNNs while cold colors the visual-related ones), when the other three DNNs are held constant. The correlations and partial correlations are based on the ROI contacts averaged response in the 0.1-0.4 sec time window. As can be seen in the figure, the pattern of correlations and partial correlations is similar to what we found in the face-selective ventral contacts (see Main text Figure 3A top and bottom).

Supplementary Figure 3 shows correlations across different time windows of the face-selective ventral contacts and the fronto-parietal visual contacts with GPT output layer. As can be seen in the Fronto-parietal contacts (right panel), the correlation of the first time window (0.1-0.4 sec) was significantly higher than all other time window, and there was no significant difference between any other pair of time windows. For the face-selective contacts (left panel), the correlation of the first time window (0.1-0.4 sec) was significantly higher than all other time window except the last one after stimulus onset (1.5-1.8 sec), which was significantly higher than the 0.7-1.0 sec and the 1-1.3 sec time windows. No other significant differences were found.

**Supplementary Figure 3:**
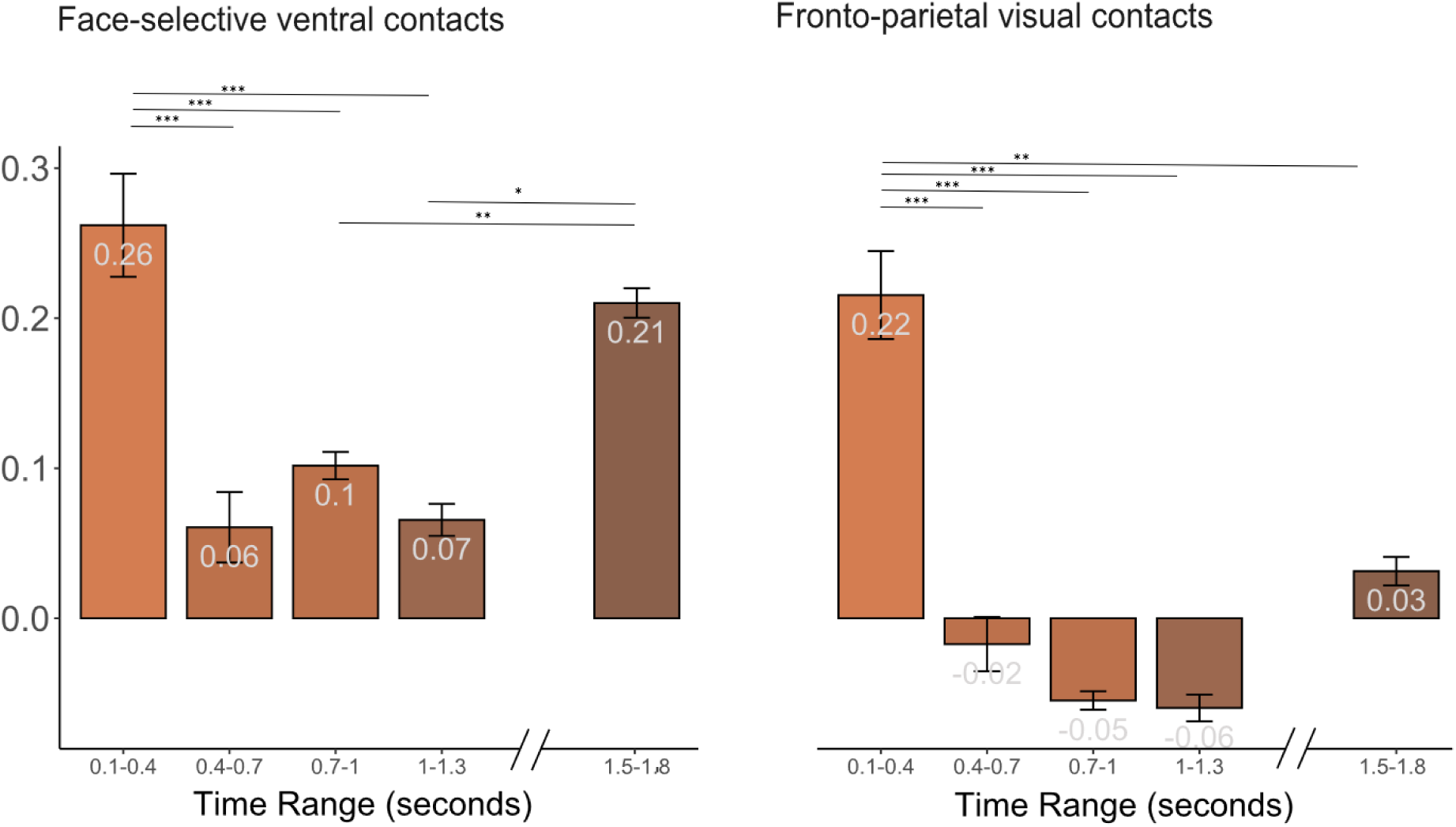
Correlations with GPT in Face selective ventral and Frontal-parietal visual contacts at different time windows: Left panel shows the correlations with the face-selective ventral contacts, and the right panel shows the correlation with the fronto-parietal visual contacts. Error bars indicate leave one participant out procedure s.e.m ∗ 𝑝_𝐹𝐷𝑅_ < 0.05, ∗∗ 𝑝_𝐹𝐷𝑅_ < 0.01, ∗∗∗ 𝑝_𝐹𝐷𝑅_ < 0.001

Supplementary Figure 4 depicts bootstrap analyses to assess the consistency of the correlations with the averaged RDM (shown in main text Figure 5): Panel A presents the average result of all possible iterations in which one repetition was removed from the analysis each time (leave one repetition out). Pannel B presents the average result of all possible iterations in which one stimulus was removed from the analysis each time (leave one stimulus out). Pannel C presents the average result of all possible iterations in which one contact was removed from the analysis each time (leave one contact out). Pannel D presents the average result of all possible iterations in which one participant was removed from the analysis each time (leave one participant out). As can be seen, the results remained fairly robust -arguing against an outlier-driven effect.

**Supplementary Figure 4:**
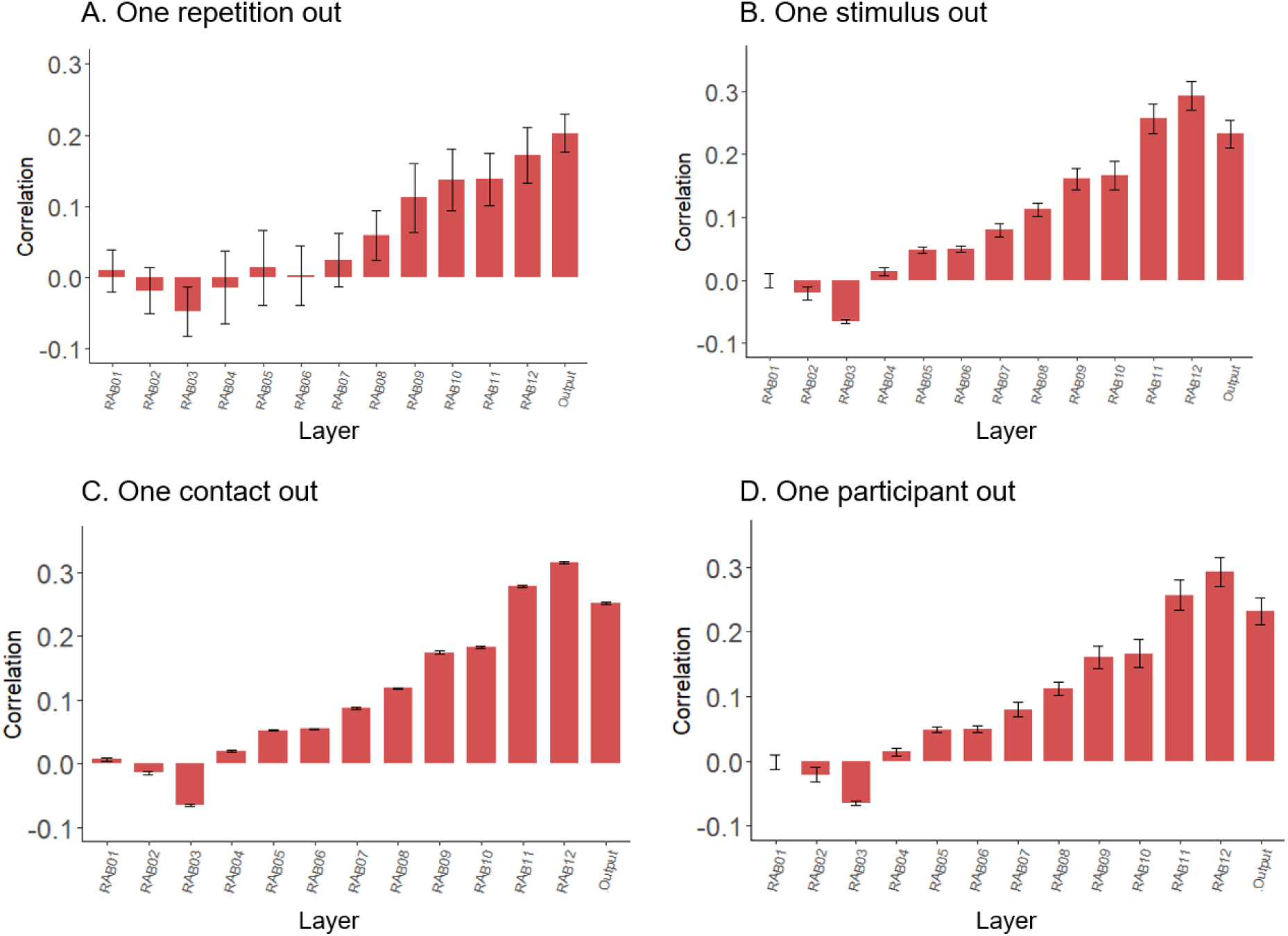
CLIP-Text’s layers pattern of correlations with Fronto-parietal visual contacts is stable for ‘leave one out‘ procedures of the following factors: The average correlation of fronto-parietal in the first time window (0.1-0.4 sec) with CLIP-Text layers for **A.** leave one repetition out. **B.** leave one stimulus out **C.** leave one contact out. **D.** leave one participant out. In all panels, the error bars represent mean standard error (se).

**Supplementary Figure 5:**
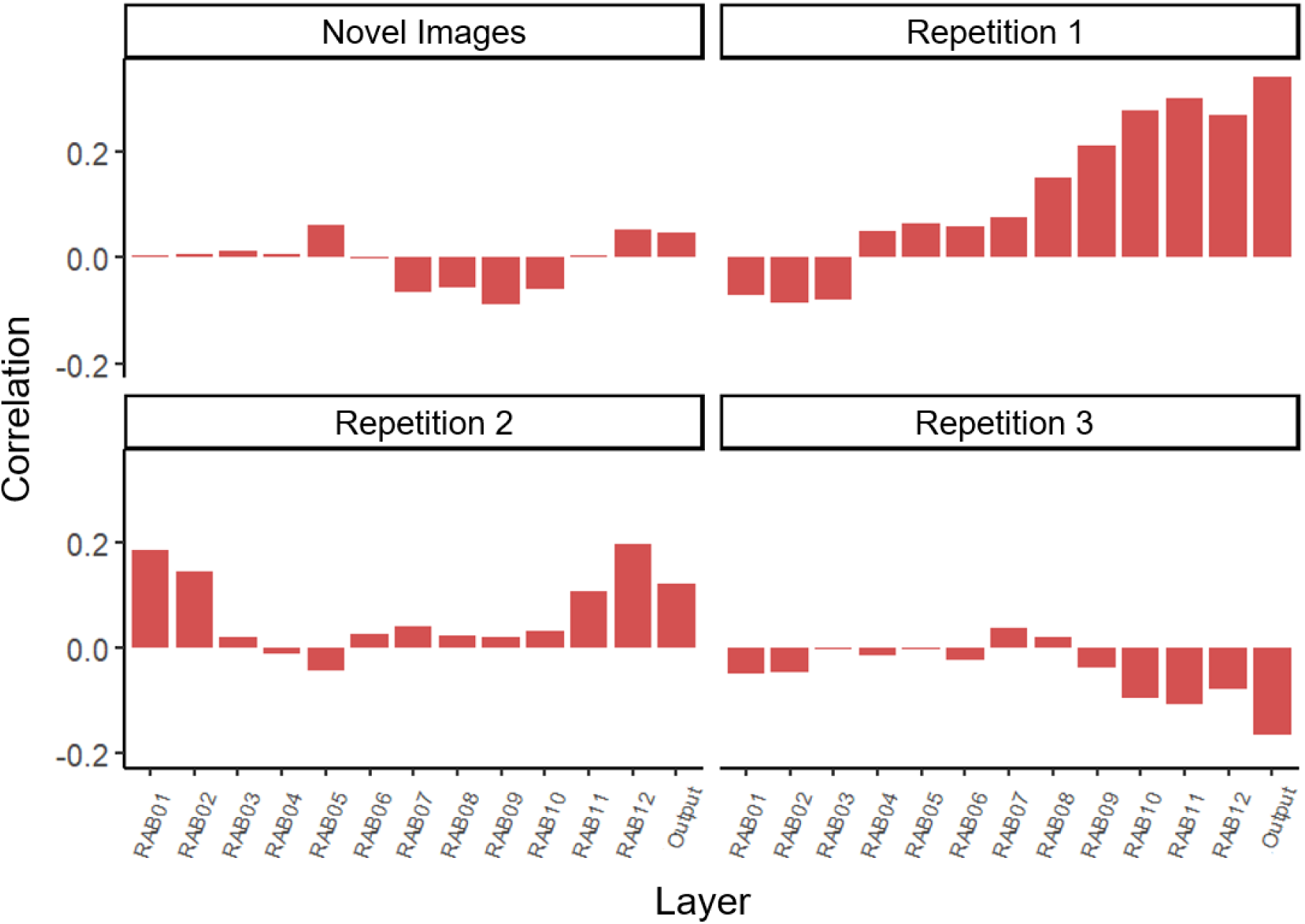
CLIP-Text’s layers pattern of correlations with Fronto-parietal contacts in each repetition separately: The correlation of fronto-parietal contacts in the first time window (0.1-0.4 sec) with CLIP-Text layers for novel images (first repetition, top-left), first repetition (top-right), second repetition (bottom-left) and third repetition (bottom-right).

To examine whether the correlation to CLIP-Text changes across different frontal subdivisions, the frontal contacts were divided into two major anatomical subdivisions: an orbitofrontal cortex (OFC) contacts group (encompassing the lateral and medial orbitofrontal cortex; n=20) and a lateral prefrontal group (including the superior frontal gyrus, middle frontal gyrus, precentral gyrus, pars orbitalis, pars triangularis and pars opercularis; n=35). Separate RDMs were constructed based on the responses of the visual contacts in each anatomical sub-division. The correlations of the RDMs of the two anatomical subdivisions with CLIP-Text. are depicted in Supplementary Figure 6. To keep as many contacts as possible we used one stimulus out instead of one participant out procedure compute the standard error of the mean.

**Supplementary Figure 6:**
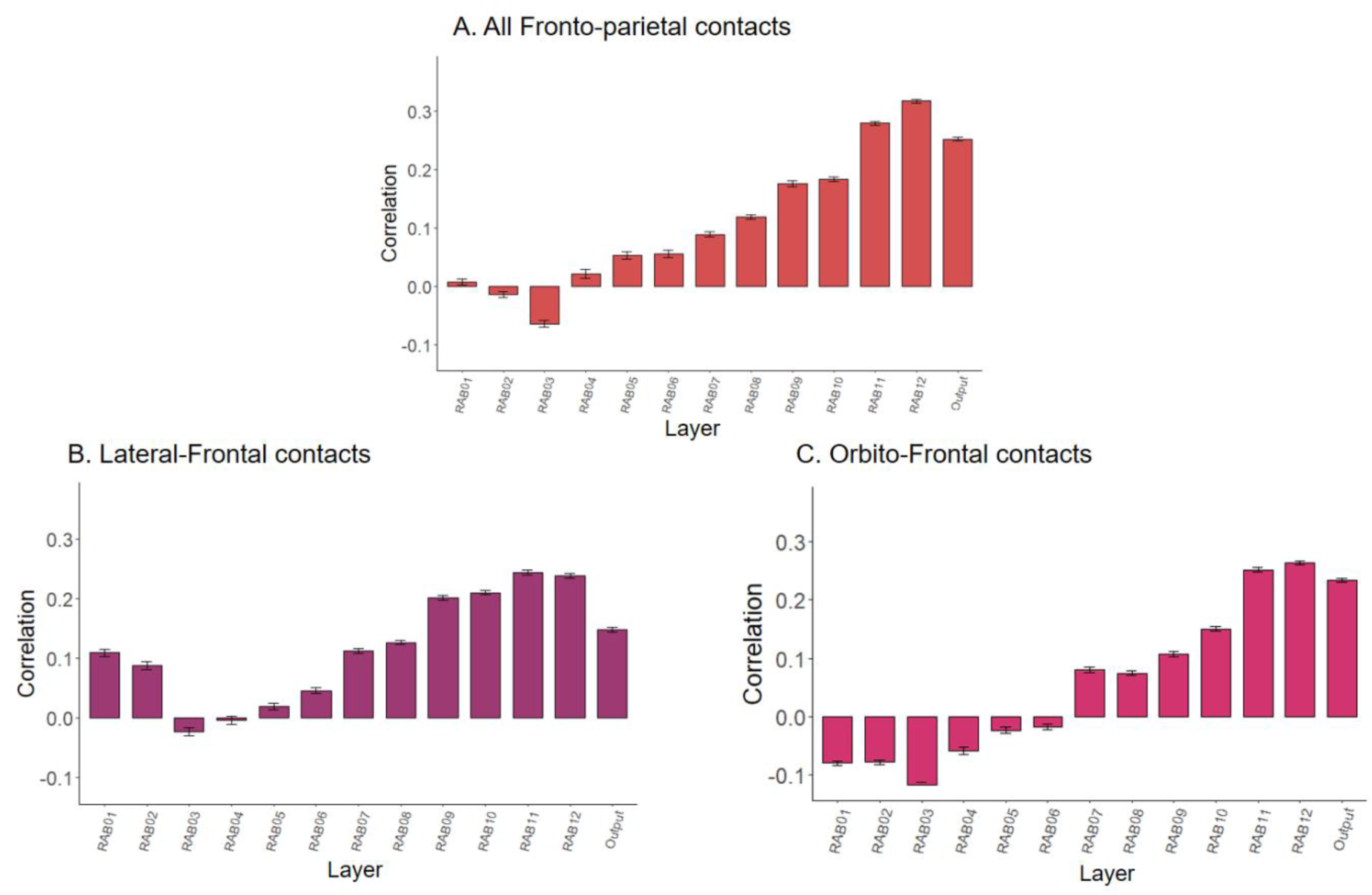
CLIP-Text’s layers pattern of correlations with sub-divisions of Fronto-parietal contacts: The correlation of fronto-parietal sub-divisions in the first time window (0.1-0.4 sec) with CLIP-Text: A. All fronto-parietal visual contacts B. Orbito-Frontal visual contacts C. Lateral-Frontal visual contacts. Error bars indicate leave one stimulus out procedure s.e.m.

**Supplementary Figure 7:**
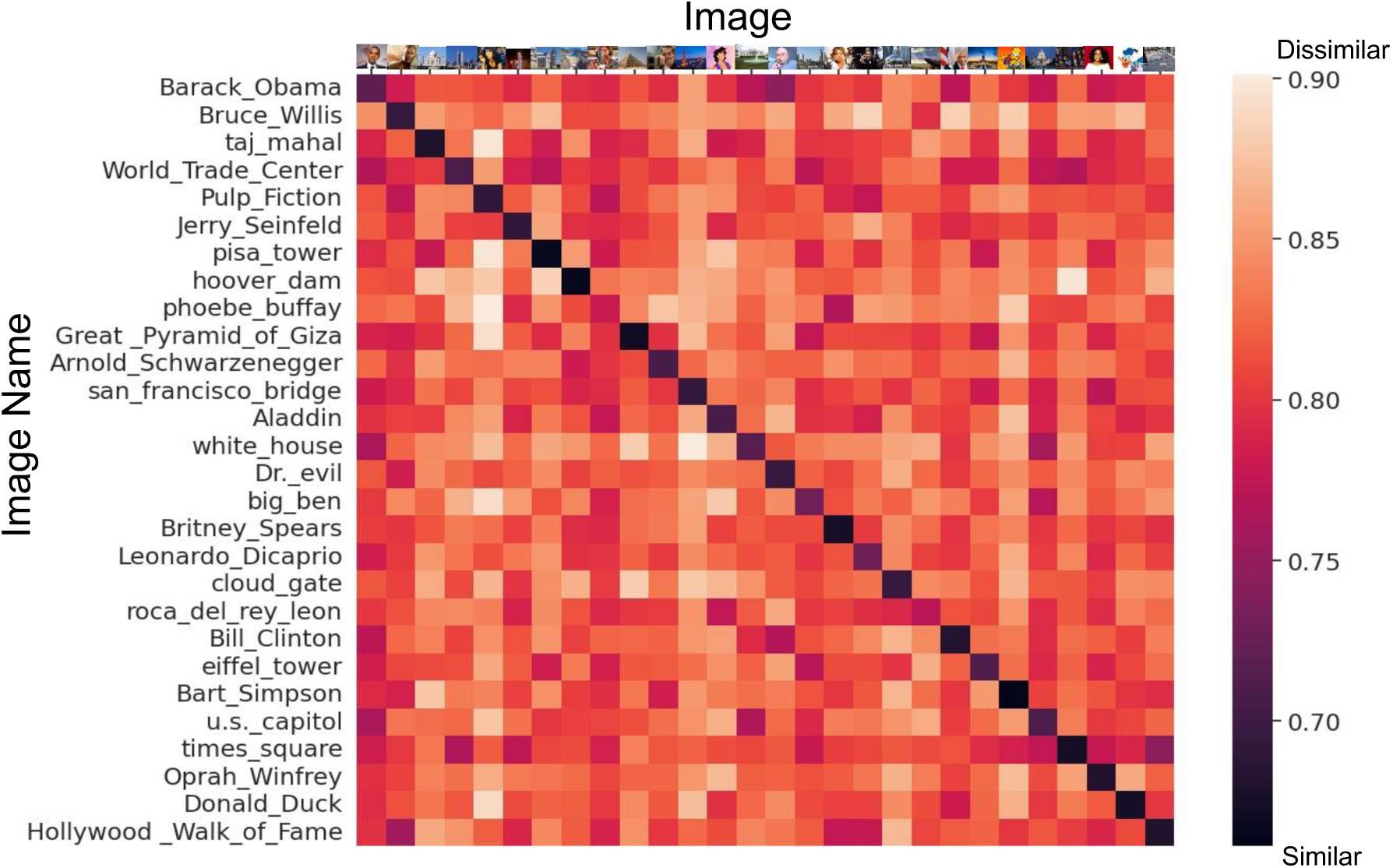
CLIP familiarity RDM: the similarity between the embeddings of each image based on CLIP-image and its name based on CLIP-text: The similarity scores between the embeddings of all images and all names of the stimuli using CLIP-Image and CLIP-Text, respectively. The distance between each image and its name was the smallest (e.g., Barack Obama’s image was closest to the text “Barack Obama” compared to any other name in the list), suggesting that CLIP’s name-based representations carry relevant semantic information.

Extracting CLIP-Text representations based on Wikipedia definitions: To determine if we missed valuable information by using only names for extracting CLIP-Text representations, we also extracted representations based on Wikipedia definitions (limited to 77 characters). We found that the correlation between CLIP-Text and CLIP-Image increased to 0.49 (compared to 0.29 based solely on names; see Figure 2B in the main text). This aligns with findings by Huh and colleagues^1^, who showed in an extensive review of vision and language DNNs that CLIP-Text representations become more similar to visual representations as the number of words in the input increases. However, CLIP-Text Wikipedia based representations showed lower correlations with brain activity (r = 0.02 with Face-selective ventral contacts and r = 0.05 with Fronto-parietal visual contacts). Therefore, we conclude that CLIP-Text’s name-based representations are more informative and suitable for predicting human brain responses.

**Supplementary Figure 8:**
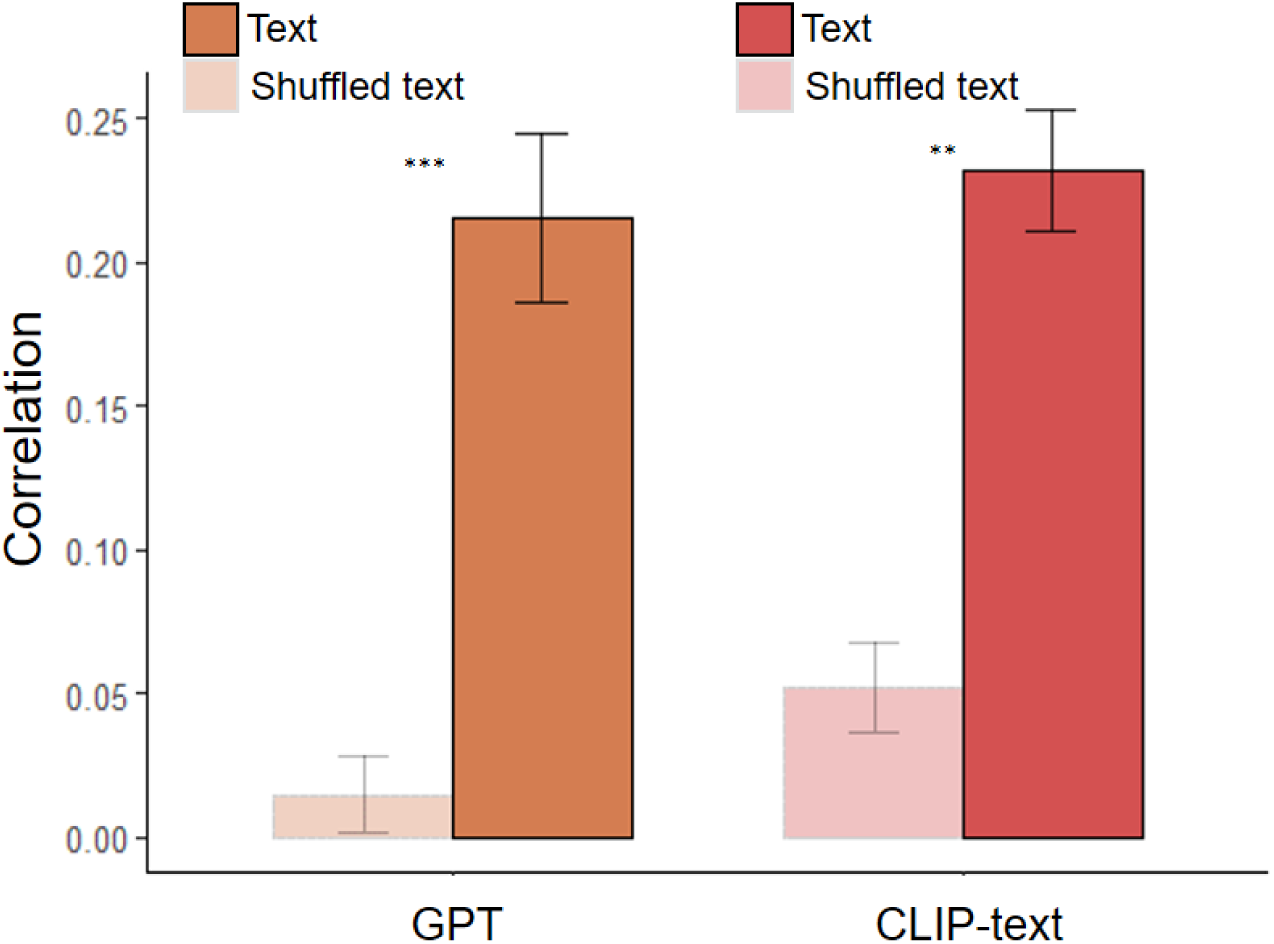
Test-based DNNs correlations with Fronto-parietal visual contacts when embeddings were based on the original text or on shuffled text: The correlations with the fronto-parietal visual contacts in the first time window (0.1-0.4 sec) with GPT output layer (orange bars) and CLIP-Text output layer (red bars), based on the original text (opaque) or shuffled text (pale). Error bars indicate leave one participant out procedure s.e.m. ∗∗ 𝑝_𝐹𝐷𝑅_ < 0.01 ∗∗∗ 𝑝_𝐹𝐷𝑅_ < 0.001. The correlation to the Text DNNs based on shuffled text input instead of the original text is drastically reduced

Supplementary Table 1 presents the text used as input for each of the text-based DNNs. The names of the identities/places featured in the images were used as CLIP-Text inputs or as the searched definition in Wikipedia. For some images, a single name could not be defined. Therefore, we used Google Image Search to determine the appropriate names. We uploaded the image and selected the first name that appeared in the search results. For instance, the image we labelled “Pulp Fiction” features the actress Uma Thurman from the well-known poster of the movie Pulp Fiction. Upon searching this image on Google Images, the top result was “Pulp Fiction,” and thus, this was the name we used.

**Supplementary Table 1:**

| Name used to extract CLIP-Text representations | Text used to extract GPT representations (Wikipedia definitions) |
| --- | --- |
| Aladdin | Aladdin is a fictional character and the titular protagonist of Walt Disney Pictures' 31st animated feature of the same name based on Aladdin, a folk tale of Middle Eastern origin. He is voiced by American actor Scott Weinger, while his singing voice is provided by Brad Kane. He also stars in the two direct-to-video sequels The Return of Jafar (1994) and Aladdin and the King of Thieves (1996), as well as the animated television series based on the film. Mena Massoud played a live-action version of the character in a live action adaptation of the 1992 film. |
| Arnold Schwarzenegger | Arnold Alois Schwarzenegger( born July 30, 1947) is an Austrian-American actor, film producer, businessman, former bodybuilder and politician who served as the 38th governor of California between 2003 and 2011. As of 2022, he is the most recent Republican governor of California. Time magazine named Schwarzenegger one of the 100 most influential people in the world in 2004 and 2007.[2][3] |
| Barack Obama | Barack Hussein Obama II (born August 4, 1961) is an American politician who served as the 44th president of the United States from 2009 to 2017. He was the first African-American president of the United States.[2] A member of the Democratic Party, he previously served as a U.S. senator from Illinois from 2005 to 2008 and as an Illinois state senator from 1997 to 2004. |
| Bart Simpson | Bart Simpson is a fictional character in the American animated television series The Simpsons and part of the Simpson family. He is voiced by Nancy Cartwright and first appeared on television in The Tracey Ullman Show short Good Night on April 19, 1987. Cartoonist Matt Groening created and designed Bart while waiting in the lobby of James L. Brooks' office. Groening had been called to pitch a series of shorts based on his comic strip, Life in Hell, but instead decided to create a new set of characters. While the rest of the characters were named after Groening's family members, Bart's name is an anagram of the word brat. After appearing on The Tracey Ullman Show for two years, the Simpson family received its own series on Fox, which debuted December 17, 1989. Bart has appeared in every Simpsons episode except Four Great Women and a Manicure. |
| Big Ben | Big Ben is the nickname for the Great Bell of the striking clock at the north end of the Palace of Westminster in London, England,[1] and the name is frequently extended to refer also to the clock and the clock tower.[2] The official name of the tower in which Big Ben is located was originally the Clock Tower, but it was renamed Elizabeth Tower in 2012 to mark the Diamond Jubilee of Elizabeth II. |
| Bill Clinton | William Jefferson Clinton ( born August 19, 1946) is an American politician who served as the 42nd president of the United States from 1993 to 2001. He previously served as governor of Arkansas from 1979 to 1981 and again from 1983 to 1992, and as attorney general of Arkansas from 1977 to 1979. A member of the Democratic Party, Clinton became known as a New Democrat, as many of his policies reflected a centrist Third Way political philosophy. He is the husband of Hillary Clinton, who was a senator from New York from 2001 to 2009, secretary of state from 2009 to 2013 and the Democratic nominee for president in the 2016 presidential election. |
| Britney Spears | Britney Jean Spears (born December 2, 1981) is an American singer, songwriter, dancer, and actress. Often referred to as the Princess of Pop, she is credited with influencing the revival of teen pop during the late 1990s and early 2000s. Regarded as a pop icon, Spears has sold nearly 150 million records worldwide, including over 70 million in the United States alone, making her one of the world's best-selling music artists.[2][3] |
| Bruce Willis | Walter Bruce Willis (born March 19, 1955) is a retired American actor. His career began on the off-Broadway stage in the 1970s.[1] He achieved fame with a leading role on the comedy-drama series Moonlighting (1985-1989) and appeared in over a hundred films, gaining widespread recognition as an action hero after his portrayal of John McClane in the Die Hard franchise (1988-2013) and other roles.[2][3] |
| Cloud Gate | Cloud Gate is a public sculpture by Indian-born British artist Anish Kapoor, that is the centerpiece of AT&T Plaza at Millennium Park in the Loop community area of Chicago, Illinois. The sculpture and AT&T Plaza are located on top of Park Grill, between the Chase Promenade and McCormick Tribune Plaza & Ice Rink. Constructed between 2004 and 2006, the sculpture is nicknamed the Bean because of its shape, a name Kapoor initially disliked, but later grew fond of. Made up of 168 stainless steel plates welded together, its highly polished exterior has no visible seams. It measures 33 by 66 by 42 feet (10 by 20 by 13 m), and weighs 110 short tons (100 t; 98 long tons). |
| Donald Duck | Donald Fauntleroy Duck[4] is a cartoon character created by The Walt Disney Company. Donald is an anthropomorphic white duck with a yellow-orange bill, legs, and feet. He typically wears a sailor shirt and cap with a bow tie. Donald is known for his semi-intelligible speech and his mischievous, temperamental, and pompous personality. Along with his friend Mickey Mouse, Donald was included in TV Guide's list of the 50 greatest cartoon characters of all time in 2002,[5] and has earned a star on the Hollywood Walk of Fame. He has appeared in more films than any other Disney character,[6] and is the most published comic book character in the world outside of the superhero genre.[7] |
| Dr. evil | Douglas Powers commonly known as Dr. Evil, is a fictional character portrayed film series. He is the main antagonist and Austin Powers and secret twin brother). He is James Bond villains, primarily Ernst Stavro Blofeld. Dr. Evil routinely hatches schemes to terrorize and take over the world, and is usually accompanied by Number Two, a goon who fronts his evil corporation Virtucon Industries, his cat Mr. Bigglesworth and his sidekick Mini-Me, a dwarf clone of himself. |
| Eiffel Tower | The Eiffel Tower is a wrought-iron lattice tower on the Champ de Mars in Paris, France. It is named after the engineer Gustave Eiffel, whose company designed and built the tower.<br>Locally nicknamed La dame de fer (French for Iron Lady), it was constructed from 1887 to 1889 as the centerpiece of the 1889 World's Fair and was initially criticised by some of France's leading artists and intellectuals for its design, but it has become a global cultural icon of France and one of the most recognisable structures in the world.[3] The Eiffel Tower is the most visited monument with an entrance fee in the world; 6.91 million people ascended it in 2015. The Tower was made a Monument historique in 1964 and named part of UNESCO World Heritage Site (Paris, Banks of the Seins) in 1991.[4] |
| Great Pyramid of Giza | The Great Pyramid of Giza is the largest Egyptian pyramid and tomb of Khufu. Built in the 26th century BC during a period of around 27 years, it is the oldest of the Seven Wonders of the Ancient World, and the only one to remain largely intact. As part of the Giza pyramid complex, it borders present-day Giza in Greater Cairo, Egypt. |
| Hollywood Walk of Fame | The Hollywood Walk of Fame is a historic landmark which consists of more than 2,700 five-pointed terrazzo and brass stars embedded in the sidewalks along 15 blocks of Hollywood Boulevard and three blocks of Vine Street in Hollywood, California. The stars are permanent public monuments to achievement in the entertainment industry, bearing the names of a mix of musicians, actors, directors, producers, musical and theatrical groups, fictional characters, and others. The Walk of Fame is administered by the Hollywood Chamber of Commerce and maintained by the self-financing Hollywood Historic Trust. It is a popular tourist destination, with an estimated 10 million annual visitors in 2010.[2] The Hollywood Chamber of Commerce holds trademark rights to the Hollywood Walk of Fame.[3] |
| Hoover Dam | Hoover Dam is a concrete arch-gravity dam in the Black Canyon of the Colorado River, on the border between the U.S. states of Nevada and Arizona. It was constructed between 1931 and 1936 during the Great Depression and was dedicated on September 30, 1935, by President Franklin D. Roosevelt. Its construction was the result of a massive effort involving thousands of workers, and cost over one hundred lives. It was referred to as Hoover Dam after President Herbert Hoover in bills passed by Congress during its construction, but was named Boulder Dam by the Roosevelt administration. The Hoover Dam name was restored by Congress in 1947. |
| Jerry Seinfeld | Jerome Allen Seinfeld (born April 29, 1954)[2] is an American stand-up comedian, actor, writer, and producer. He is best known for playing a semi-fictionalized version of himself in the sitcom Seinfeld, which he created and wrote with Larry David. The show aired on NBC from 1989 until 1998, becoming one of the most acclaimed and popular American sitcoms of all time. As a stand-up comedian, Seinfeld specializes in observational comedy. In 2004, Comedy Central named him as the 12th-greatest stand-up comedian of all time |
| Leonardo DiCaprio | Leonardo Wilhelm DiCaprio (born November 11, 1974) is an American actor and film producer. Known for his work in biopics and period films, he is the recipient of numerous accolades, including an Academy Award, a British Academy Film Award, and three Golden Globe Awards. As of 2019, his films have grossed over \$7.2 billion worldwide, and he has been placed eight times in annual rankings of the world's highest-paid actors. |
| Oprah Winfrey | Oprah Gail Winfrey ( January 29, 1954) is an American talk show host, television producer, actress, author, and philanthropist. She is best known for her talk show, The Oprah Winfrey Show, broadcast from Chicago, which was the highest-rated television program of its kind in history and ran in national syndication for 25 years, from 1986 to 2011.[3][better source needed][4] Dubbed the Queen of All Media,[5] she was the richest African-American of the 20th century,[6][7] was once the world's only black billionaire,[8] and the greatest black philanthropist in U.S. history.[9][10] By 2007, she was sometimes ranked as the most influential woman in the world.[11][12] |
| Phoebe Buffay | Phoebe Buffay is one of the six main characters from the American sitcom Friends. She is created by David Crane and Marta Kauffman and portrayed by actress Lisa Kudrow.<br>In the series' universe, Phoebe was born on February 16 and is the daughter of Frank and Lily Buffay. Her biological mother's name was Phoebe Abbott, whom she was named after. Phoebe has a twin sister, Ursula, a waitress who is also portrayed by Kudrow. Phoebe can speak several languages, including French and Italian. She appeared in all of the show's 236 episodes. |
| Pisa Tower | The Leaning Tower of Pisa (Italian: torre pendente di Pisa), or simply, the Tower of Pisa, is the campanile, or freestanding bell tower, of Pisa Cathedral. It is known for its nearly four-degree lean, the result of an unstable foundation. The tower is one of three structures in the Pisa's Cathedral Square (Piazza del Duomo), which includes the cathedral and Pisa Baptistry. |
| Pulp Fiction | Pulp Fiction is a 1994 American black comedy crime film written and directed by Quentin Tarantino, who conceived it with Roger Avary.[4] Starring John Travolta, Samuel L. Jackson, Bruce Willis, Tim Roth, Ving Rhames, and Uma Thurman, it tells several stories of crime in Los Angeles. The title refers to the pulp magazines and hardboiled crime novels popular during the mid-20th century, known for their graphic violence and punchy dialogue. |
| The Lion King | The Lion King is a 1994 American animated musical drama film[4] produced by Walt Disney Feature Animation and released by Walt Disney Pictures. It is the 32nd Disney animated feature film and the fifth animated film produced during a period known as the Disney Renaissance. The Lion King was directed by Roger Allers and Rob Minkoff (in their feature directorial debuts) and produced by Don Hahn, from a screenplay by Irene Mecchi, Jonathan Roberts, and Linda Woolverton. Its original songs were written by composer Elton John and lyricist Tim Rice, with a score by Hans Zimmer. The film features an ensemble voice cast that includes Matthew Broderick, James Earl Jones, Jeremy Irons, Jonathan Taylor Thomas, Moira Kelly, Nathan Lane, Ernie Sabella, Rowan Atkinson, Robert Guillaume, Madge Sinclair (in her last film role), Whoopi Goldberg, Cheech Marin, and Jim Cummings. |
| San francisco bridge | The Golden Gate Bridge is a suspension bridge spanning the Golden Gate, the one-mile-wide (1.6 km) strait connecting San Francisco Bay and the Pacific Ocean. The structure links the U.S. city of San Francisco, California the northern tip of the San Francisco Peninsula to Marin County, carrying both U.S. Route 101 and California State Route 1 across the strait. It also carries pedestrian and bicycle traffic, and is designated as part of U.S. Bicycle Route 95. Being declared one of the Wonders of the Modern World by the American Society of Civil Engineers,[7] the bridge is one of the most internationally recognized symbols of San Francisco and California. It was initially designed by engineer Joseph Strauss in 1917. |
| Taj Mahal | The Taj Mahal is an ivory-white marble mausoleum on the right bank of the river Yamuna in the Indian city of Agra. It was commissioned in 1632 by the Mughal emperor Shah Jahan (r. 1628 1658) to house the tomb of his favourite wife, Mumtaz Mahal; it also houses the tomb of Shah Jahan himself. The tomb is the centrepiece of a 17-hectare (42-acre) complex, which includes a mosque and a guest house, and is set in formal gardens bounded on three sides by a crenellated wall. |
| Times square | Times Square is a major commercial intersection, tourist destination, entertainment center, and neighborhood in Midtown Manhattan, New York City. It is formed by the junction of Broadway, Seventh Avenue and 42nd Street. Together with adjacent Duffy Square, Times Square is a bowtie-shaped space five blocks long between 42nd and 47th Streets. |
| U.S. capitol | The United States Capitol, often called The Capitol or the Capitol Building, is the meeting place of the United States Congress and the seat of the legislative branch of the U.S. federal government. It is located on Capitol Hill at the eastern end of the National Mall in Washington, D.C. Though no longer at the geographic center of the federal district, the Capitol forms the origin point for the district's street-numbering system and the district's four quadrants. |
| White House | The White House is the official residence and workplace of the president of the United States. It is located at 1600 Pennsylvania Avenue NW in Washington, D.C., and has been the residence of every U.S. president since John Adams in 1800. The term White House is often used as a metonym for the president and his advisers. |
| World Trade Center | The original World Trade Center (WTC) was a large complex of seven buildings in the Financial District of Lower Manhattan, New York City, United States. It opened on April 4, 1973, and was destroyed in 2001 during the September 11 attacks. At the time of their completion, the Twin Towers—the original 1 World Trade Center (the North Tower) at 1,368 feet (417 m); and 2 World Trade Center (the South Tower) at 1,362 feet (415.1 m)—were the tallest buildings in the world. Other buildings in the complex included the Marriott World Trade Center (3 WTC), 4 WTC, 5 WTC, 6 WTC, and 7 WTC. The complex contained 13,400,000 square feet (1,240,000 m<sup>2</sup>) of office space. |

